# Aster-dependent non-vesicular transport facilitates dietary cholesterol uptake

**DOI:** 10.1101/2023.07.07.548168

**Authors:** Alessandra Ferrari, Emily Whang, Xu Xiao, John P. Kennelly, Beatriz Romartinez-Alonso, Julia J. Mack, Thomas Weston, Kai Chen, Youngjae Kim, Marcus J. Tol, Lara Bideyan, Alexander Nguyen, Yajing Gao, Liujuan Cui, Alexander H. Bedard, Jaspreet Sandhu, Stephen D. Lee, Louise Fairall, Kevin J. Williams, Wenxin Song, Priscilla Munguia, Robert A. Russell, Martin G. Martin, Michael E. Jung, Haibo Jiang, John W.R. Schwabe, Stephen G. Young, Peter Tontonoz

## Abstract

Intestinal cholesterol absorption is an important contributor to systemic cholesterol homeostasis. Niemann-Pick C1 Like 1 (NPC1L1), the target of the drug ezetimibe (EZ), assists in the initial step of dietary cholesterol uptake. However, how cholesterol moves downstream of NPC1L1 is unknown. Here we show that Aster-B and Aster-C are critical for non-vesicular cholesterol movement in enterocytes, bridging NPC1L1 at the plasma membrane (PM) and ACAT2 in the endoplasmic reticulum (ER). Loss of NPC1L1 diminishes accessible PM cholesterol in enterocytes and abolishes Aster recruitment to the intestinal brush border. Enterocytes lacking Asters accumulate cholesterol at the PM and display evidence of ER cholesterol depletion, including decreased cholesterol ester stores and activation of the SREBP-2 transcriptional pathway. Aster-deficient mice have impaired cholesterol absorption and are protected against diet-induced hypercholesterolemia. Finally, we show that the Aster pathway can be targeted with a small molecule inhibitor to manipulate dietary cholesterol uptake. These findings identify the Aster pathway as a physiologically important and pharmacologically tractable node in dietary lipid absorption.

**One-Sentence Summary:** Identification of a targetable pathway for regulation of dietary cholesterol absorption

## Main text

The intestine regulates systemic lipid homeostasis by gating dietary cholesterol intake (*1*). Exogenous cholesterol absorbed at the apical PM of enterocytes is trafficked to the ER, where it is esterified by acyl CoA:cholesterol acyltransferase 2 (ACAT2) for chylomicron packaging (*2*). Niemann-Pick C1-Like 1 (NPC1L1), an important mediator of intestinal cholesterol uptake (*3, 4*) and a target of the drug ezetimibe (EZ) (*5, 6*), has been shown to facilitate cholesterol movement into the enterocyte PM (*7*). Statins are effective in lowering LDL-cholesterol, but adding EZ to statin treatment further reduces cholesterol levels and cardiovascular events compared to statin therapy alone in humans (*8*), indicating the intestine remains a potential therapeutic target for the regulation of cholesterol homeostasis.

EZ was initially discovered as a weak inhibitor of ACAT2 (*9*). Its profound effects on cholesterol absorption led to its FDA approval in 2002, even before the discovery that it inhibits NPC1L1. Genetic ablation of NPC1L1 or pharmacologic inhibition by EZ impairs cholesterol absorption (*3*), while ACAT2 deficiency reduces cholesterol ester production required for chylomicron assembly (*10–13*). Treatment with EZ or ablating ACAT2 expression protects mice from diet-induced hypercholesterolemia and atherosclerosis (*1, 14–16*). Despite the importance of the NPC1L1-ACAT2 axis for cholesterol homeostasis, how cholesterol deposited by NPC1L1 at the PM reaches the ER for esterification by ACAT2 is unknown. Elucidating this pathway could establish an additional route by which cholesterol absorption may be targeted pharmacologically.

The mechanism of NPC1L1 action has been studied intensively (*17–24*). Recent structural analyses have revealed that the N-terminal domain (NTD) of NPC1L1 possesses a cavity that accommodates cholesterol (*7, 25, 26*). After cholesterol binds, the NTD assumes a new configuration in which the cholesterol binding pocket faces a passageway that connects to the sterol-sensing domain (SSD) of NPC1L1. Cholesterol moves through this channel to reach the SSD and diffuses into the lipid bilayer. EZ binds within the channel, blocking cholesterol deposition in the PM.

Aster proteins (Aster-A, -B, and -C; encoded by *Gramd1a, Gramd1b,* and *Gramd1c*) bind to cholesterol and facilitate its movement between membranes (*27–29*). Members of this family have a central ASTER domain with a cholesterol-binding pocket that is flanked by a GRAM domain at the N-terminus and by an ER transmembrane domain at the C-terminus. The GRAM domain binds the PM in response to cholesterol loading, allowing the ASTER domain to transfer cholesterol down a concentration gradient from the PM to the ER. In mice, loss of Aster-B results in the depletion of cholesterol esters in the adrenal cortex and impairs corticosteroid synthesis (*27*). However, the role of the Aster pathway in systemic sterol homeostasis is unknown.

Dietary cholesterol and fatty acids are absorbed by enterocytes and packaged into chylomicrons, which are released into the lymphatics and ultimately reach the systemic circulation (*30*). Most of the cholesterol in chylomicrons is esterified, and cholesterol ester (CE) is necessary for chylomicron packaging. Free cholesterol deposited into the inner leaflet of the apical PM by NPC1L1 must first move to the ER, where it is esterified by ACAT2 for chylomicron assembly. ACAT2 deficiency reduces the efficiency of cholesterol absorption and its incorporation into chylomicrons (*11, 12, 15*). How cholesterol is delivered to the enterocyte ER for esterification remains to be defined.

Here we show that Aster-B and -C cooperate with NPC1L1 to deliver dietary cholesterol from the intestinal lumen to the enterocyte ER. Asters are recruited to the enterocyte PM upon cholesterol loading, and loss of Aster expression impairs cholesterol delivery to the ER and CE production. Mice lacking Asters in the intestine exhibit reduced cholesterol absorption and are protected from diet-induced hypercholesterolemia. Treatment of mice with a small-molecule Aster inhibitor reduces the systemic absorption of dietary cholesterol. Our findings support a model in which NPC1L1 enriches the enterocyte PM with cholesterol, followed by Aster-mediated cholesterol trafficking to the ER. These findings identify intestinal Asters as a potential target for the control of cholesterol homeostasis.

## Results

### Aster protein expression in the small intestine is regulated by Liver X Receptors

Transcripts for Aster-A, -B, and -C are expressed in the small intestine (SI) (Fig.1A), but those for Aster-B and -C are most abundant in intestinal epithelial cells (IEC) (Fig. S1B). RNA-seq from proximal jejunal scrapings of C57BL6/J mice confirmed robust expression of *Gramd1b* and *Gramd1c* and low expression of *Gramd1a* (Fig. S1A-C). The most abundant *Gramd1c* transcript in the SI encodes a truncated Aster-C lacking the N-terminal GRAM domain. RNA- seq also revealed the presence of two transcripts for *Gramd1b* in the intestine; one is the same as that identified previously in macrophages (*27*), while the other is longer and has an extended N-terminal region upstream of the GRAM domain (Fig. S1A). An intestine-specific promoter region in the *Gramd1b* locus (chr9: 40465470-40465756) defines the longer variant. We found that Aster-B was expressed along the entire length of the SI (duodenum to the ileum) (Fig. 1B). In Caco-2 cells, transcripts for Aster-B and -C (but not Aster-A) were induced upon differentiation (Fig. S1C). Liver X receptor (LXR) transcription factors are key regulators of cholesterol metabolism, including in the intestine (*31, 32*). Treatment of WT mice with the LXR agonist GW3965 induced the expression of all three Asters in SI (Fig. S1D), consistent with a role for the Aster pathway in intestinal cholesterol flux.

**Fig. 1:**
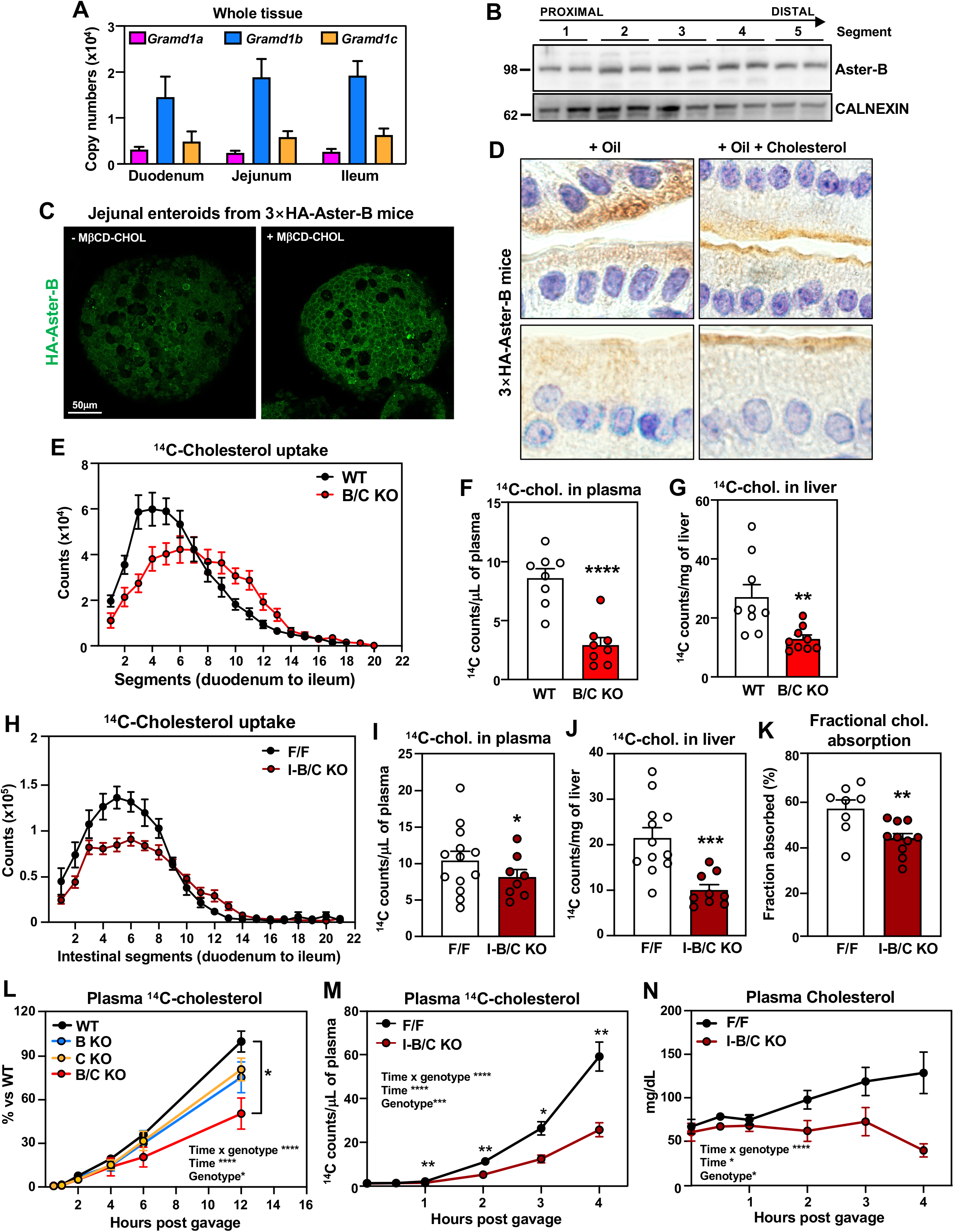
Aster proteins modulate dietary cholesterol uptake in the small intestine. (**A**) Absolute quantification of mRNA of *Gramd1a*, *Gramd1b*, and *Gramd1c* in homogenized proximal jejunum; (**B**) Western blot analysis of expression of Aster-B along the small intestine in C57BL/6J male mice (*n* = 2); (**C**) Immunofluorescence microscopy of HA-Aster-B in intestinal organoids from 3×HA-Aster-B mice during sterol deprivation (left) or loading with MβCD- cholesterol (right); (**D**) Immunohistochemistry of HA-Aster-B in small intestines from 3×HA- Aster-B mice after a gastric gavage with corn oil or corn oil with cholesterol ; (**E**) Distribution of radioactivity in intestinal segments of female WT and B/C KO mice after oral gavage with olive oil containing [^14^C]cholesterol for 2 h (*n = 9*/group); (**F**) Radioactivity in plasma of mice described in E; (**G**) Radioactivity in liver of mice described in E; (**H**) Distribution of radioactivity in intestinal segments of female Aster-B/C^fl/fl^ (F/F) and littermates Aster-B/C^fl/fl^ Villin-CreERT2 (I-B/C KO) mice after an oral challenge of olive oil containing [^14^C]cholesterol for 2 h (*n = 9*/group); (**I**) Radioactivity in plasma of mice described in H; (**J**) Radioactivity in liver of mice described in H; (**K**) *In vivo* cholesterol absorption measured by fecal dual-isotope ratio method (*n = 8–10*/group); (**L**) Kinetics of radioactivity in plasma of female WT (*n = 9*), Aster-B global knockout (B KO) (*n = 5*), Aster-C global knockout (C KO) (*n = 5*), and Aster- B/C global knockout (B/C KO) (*n = 5*) mice after an oral challenge of olive oil containing [^14^C]cholesterol; (**M**) Kinetics of radioactivity in plasma of female F/F and littermates I-B/C KO mic after injection of Poloxamer-407 and an oral challenge of olive oil containing [^14^C]cholesterol; (**N**) Kinetics of total cholesterol in mice described in M. Data are expressed as mean ± SEM. Statistical analysis: for panels F, G, I, J, K, unpaired *t* test; *p < 0.05, **p < 0.01, ***p < 0.001, ****p < 0.0001; for panel L, 2-way ANOVA with Tukey’s as multiple comparisons test; for panels M and N, 2-way ANOVA with Sidak’s as multiple comparisons test. *p < 0.05, **p < 0.01, ***p < 0.001, ****p < 0.0001.

### Aster proteins modulate cholesterol uptake in the small intestine

To visualize endogenous Aster-B movement in IECs, we used CRISPR-Cas9 editing to insert a 3×HA tag into the mouse *Gramd1b* locus (Fig. S1E and F). Immunohistochemistry of intestinal tissue revealed that HA-Aster-B was enriched in the villi of the jejunum with lower expression in the crypts (S1G). To study Aster localization in a culture model, we derived enteroids from intestinal crypts of HA-Aster-B mice. Confocal imaging demonstrated that Aster-B was recruited to the PM in response to loading with methyl-β-cyclodextrin (MbCD)– cholesterol (Fig. 1C). Accordingly, immunohistochemistry of the small intestine showed a greater distribution of HA-Aster-B at the brush border of enterocytes 1 h after intra-gastric gavage with cholesterol in corn oil versus corn oil alone (Fig. 1D). These observations implicate Aster proteins in the uptake of dietary cholesterol.

To explore the functions of Asters in intestinal physiology, we generated global knockout mice for Aster-B (B-KO), Aster-C (C-KO), or both (B/C-KO) (Fig. S2A and B). Deletion of Asters was verified in jejunal scrapings (Fig. S2C, D, and E). The SI from single-KO and B/C- KO mice had no obvious histological abnormalities (Fig. S2F), and body weight and intestinal length were similar across genotypes (Fig. S2G). To evaluate the role of Asters in cholesterol uptake, we administered [^14^C]cholesterol by gastric gavage and then measured radioactivity in SI biopsies 2 h later. No differences in [^14^C]cholesterol absorption were detected in single-KO mice (Fig. S2 H-M), but we observed markedly reduced absorption in the proximal intestine of B/C-KO mice (Fig. 1E). B/C-KO mice also had reduced amounts of [^14^C]cholesterol in the plasma and liver 2 h after gavage (Fig. 1, F and G).

To confirm that this impairment in cholesterol absorption resulted from Aster deficiency in enterocytes, we also generated mice with tamoxifen-inducible, intestine-specific deletion of Aster-B (I-B-KO), Aster-C (I-C-KO), or both (I-B/C-KO) by intercrossing double “floxed” mice with Villin-Cre^ERT2^ transgenic mice (Fig. S3A-F). Immunohistochemistry staining for Olfm4, a stem cell marker, indicated that I-B/C KO mice displayed normal crypt cell proliferation (Fig. S3G). No difference in body weight or SI length was observed in I-B/C KO mice (Fig. S3H). Thus, loss of Aster expression does not appear to compromise intestinal development.

Cholesterol uptake was similar in I-B KO and I-C-KO mice and littermate controls (Fig. S3I-N). However, I-B/C-KO mice had reduced [^14^C]cholesterol absorption (comparable to that observed in global B/C-KO mice) (Fig. 1H-J). Fractional cholesterol absorption, assessed by fecal dual isotope labeling, confirmed impaired cholesterol absorption in I-B/C-KO mice (Fig. 1K). By contrast, there were only minor reductions in ^14^C counts in intestinal segments, plasma, and liver of I-B/C-KO mice after an oral gavage of [^14^C]triolein (Fig. S4A-D). Glucose absorption was not impacted in I-B/C-KO mice (Fig. S4E). Thus, loss of Aster function in enterocytes selectively impairs cholesterol absorption.

Next, we followed the appearance of orally administered [^14^C]cholesterol in the plasma of Aster-KO mice over 12 h. There was a marked reduction in ^14^C counts in the plasma of B/C- KO mice, while B-KO and C-KO mice exhibited an intermediate phenotype (Fig. 1L). We also injected mice with the lipoprotein lipase inhibitor Poloxamer-407, gavaged them with [^14^C]cholesterol, and collected plasma up to 4 h post gavage. The appearance of [^14^C]cholesterol in the plasma over time, as well as total plasma cholesterol levels, were reduced in I-B/C-KO mice (Fig. 1M and 1N).

### Non-vesicular cholesterol transport promotes cholesterol ester production for chylomicron assembly

Adequate CE is required for chylomicron production by the intestine. We hypothesized that Aster-mediated non-vesicular transport from the PM to the ER facilitates cholesterol esterification and subsequent incorporation into chylomicrons. We used nanoscale secondary ion mass spectrometry (NanoSIMS) imaging to visualize cholesterol uptake and intracellular distribution by enterocytes (*33, 34*). WT and B/C KO mice received an intra-gastric gavage of [^13^C]fatty acids and [^2^H]cholesterol in olive oil. Biopsies from the duodenum were harvested 2 h later. Backscattered electron images of duodenal sections verified the integrity of intestinal villi (Fig. 2A and Fig. S5A). NanoSIMS images of the same sections (Fig. 2A, Fig. S5A) revealed reduced amounts of ^2^H in medial and distal segments of the duodenum in B/C-KO mice. Quantification of ^13^C^−^ and ^2^H^−^ secondary ions confirmed reduced cholesterol uptake by enterocytes lacking Asters (Fig. 2B, Fig. S5B).

**Fig. 2:**
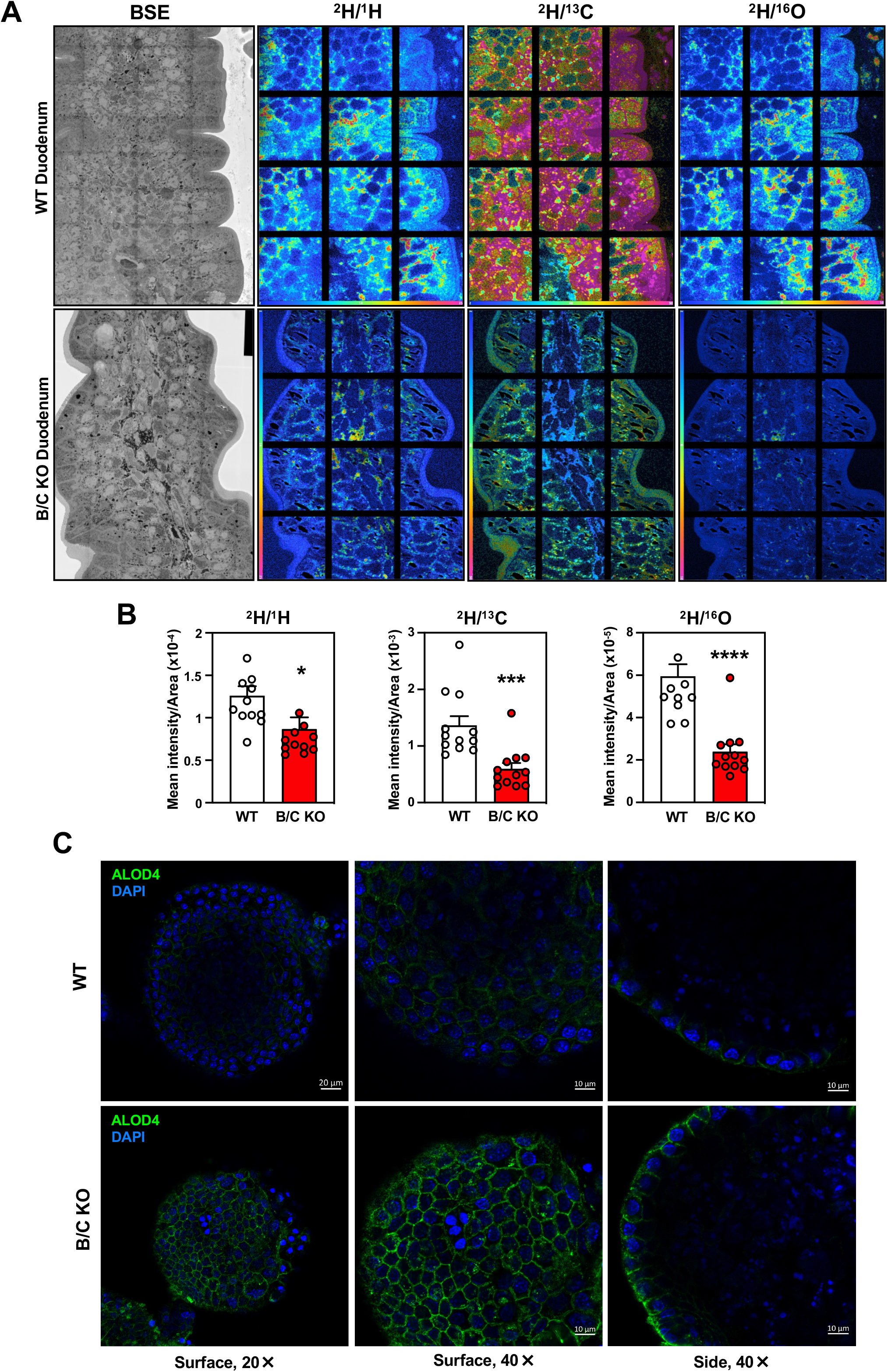
Deletion of Asters reduces cholesterol internalization from the plasma membrane in enterocytes. (**A**) Backscattered electron images and NanoSIMS images a mid-duodenum villus from WT and B/C KO mice; (**B**) Quantification of the ^2^H^−^ secondary ion signal, normalized to the ^1^H^−^, ^13^C^−^, and ^16^O^−^ signals.; (**C**) ALOD4 imaging of murine enteroids from WT and B/C KO mice. Data are expressed as mean ± SEM. Statistical analysis: unpaired *t* test; *p < 0.05, ***p < 0.001, ****p < 0.0001

Previous studies showed that Asters in macrophages (*35*) and hepatocytes (*36*) transport the accessible pool of PM cholesterol (*37, 38*). To prove that Asters transfer accessible cholesterol from PM to ER in intestinal epithelial cells, we stained WT and Aster-B/C KO enteroids with ALOD4, a bacterial peptide that selectively binds to accessible cholesterol.

Confocal microscopy revealed enhanced ALOD4 staining at the PM of Aster-B/C KO enteroids (Fig. 2C), indicating that genetic ablation of Asters in intestinal cells reduced movement of PM cholesterol to ER. Accordingly, lipidomic analysis of jejunal scrapings revealed reduced CE in B/C-KO mice after feeding (Fig. 3A). Other lipid species were not affected by Aster deficiency (Fig. S6A). Consistent with these findings, ^14^C-labeled CE accumulation was reduced in the proximal jejunum of Aster-deficient mice 2 h after oral administration of [^14^C]cholesterol in olive oil (Fig. 3B). The level of free [^14^C]cholesterol was similar in Aster-deficient mice and controls (Fig. S6B). SREBP-2 target gene expression was higher in global B/C-KO than in WT mice, both in fasted (Fig. 3C) and refed (Fig. 3D) states, consistent with a reduction in cholesterol delivery to the ER. Similar gene-expression findings were observed in jejunal scrapings of I-B/C-KO mice (Fig. 3E). These changes correlated with an increase in protein expression of SREBP-2 targets in duodenal scrapings of B/C-KO mice (Fig. 3F). We also observed upregulation of *Srebf2* and its targets in jejunal scrapings in B/C-KO mice that were fed a high-cholesterol (HC) diet (1.25% cholesterol) (Fig. S6C). Thus, even in the setting of increased amounts of dietary cholesterol, cholesterol synthesis was activated in B/C-KO mice. We did not observe changes in SREBP-2 pathway gene expression in the jejunum of global Aster-B and Aster-C single-KO mice (Fig. S6, D and E) or intestinal-specific single-KO mice (Fig. S6, F and G).

**Fig. 3:**
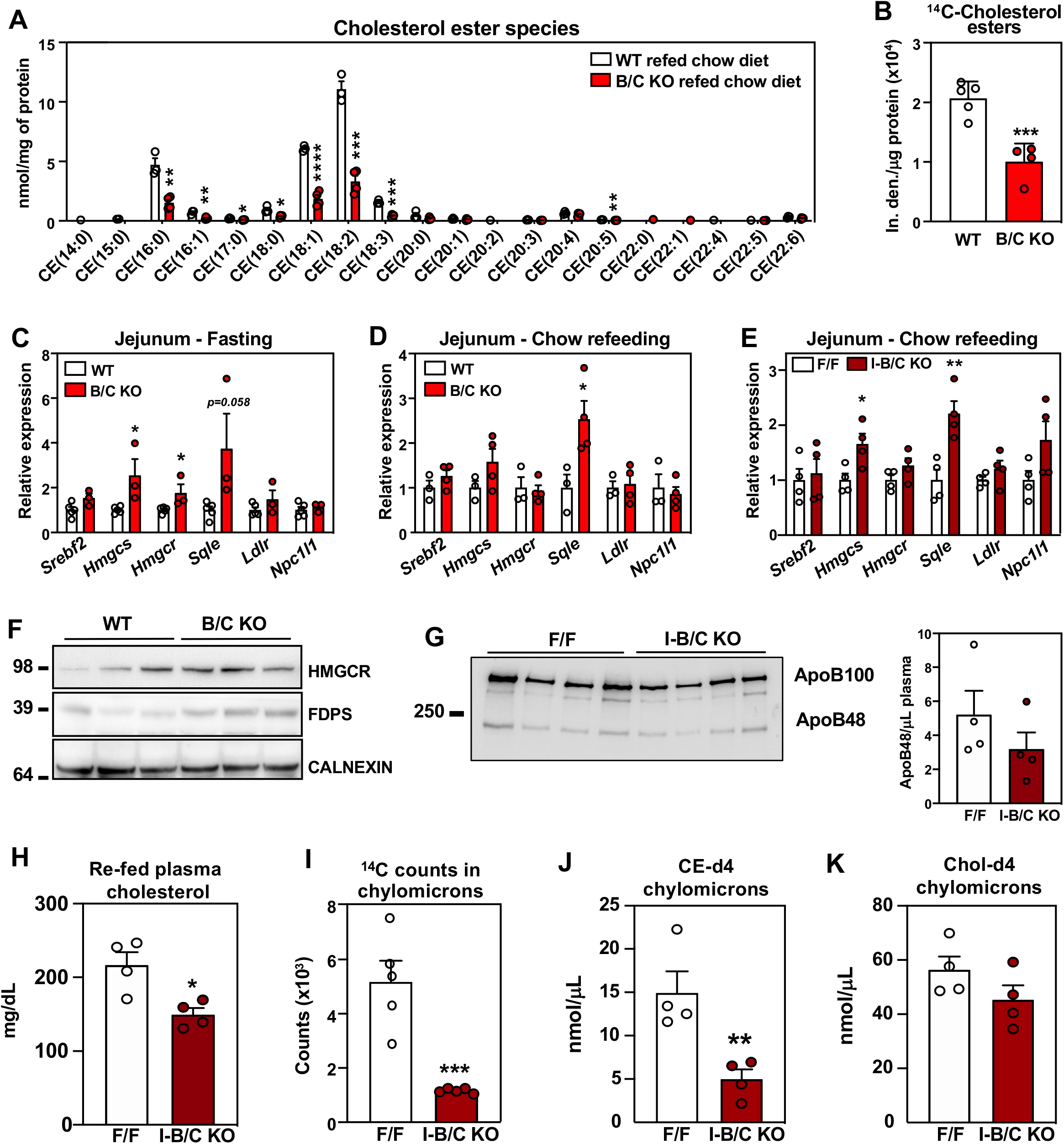
Loss of Asters in intestine impairs cholesterol transfer to ER and chylomicron packaging. (**A**) Cholesterol ester quantification by mass spectrometry in proximal jejunum from WT (n = 3) and B/C KO *(n = 4*) 2 h after refeeding a chow diet; (**B**) Quantification of ^14^C-labeled cholesterol esters isolated from proximal jejunum scrapings of WT (*n = 5*) and B/C KO (*n = 4*) mice 2 h after oral gavage of [^14^C]cholesterol; (**C**) Gene expression from distal jejunum scrapings of WT (*n = 5*) and B/C KO (*n = 3*) mice after 4 h of fasting; (**D**) Gene expression from distal jejunum scrapings of WT (*n = 5*) and B/C KO (*n = 3*) mice 2 h after refeeding a chow diet; (**E**) Gene expression from distal jejunum scrapings of F/F (*n = 4*) and I-B/C KO (*n = 4*) mice 2 h after refeeding with chow diet; (**F**) Western blot analysis of duodenum scrapings of WT and B/C KO (*n = 3*) mice 2 h after refeeding a chow diet; (**G**) Western blot analysis and quantification of plasma from F/F (*n* = 4) and I-B/C KO (*n* = 4) mice fed for 21 days a high cholesterol (1.25%) diet, 2 h post refeeding (**H**) Plasma cholesterol of mice described in G; (**I**) Quantification of ^14^C- counts in chylomicrons isolated from plasma of F/F (n=5) and I-B/C KO (n=5) 3 h after treatment with Poloxamer-407 and oral gavage of [^14^C]cholesterol; (**J**), (**K**) Quantification of deuterated (-d4) free cholesterol and CE in chylomicrons isolated from plasma of F/F (*n* = 4) and I-B/C KO (*n* = 4) 3.5 h after treatment with Poloxamer-407 and oral gavage with cholesterol-d4. Data are expressed as mean ± SEM. Statistical analysis: unpaired *t* test, *p < 0.05, **p < 0.01, ***p < 0.001, ****p < 0.0001.

Plasma levels of ApoB48 after refeeding with a HC diet were not different (Fig. 3G), despite lower plasma cholesterol levels in I-B/C KO mice (Fig. 3H). Plasma triglycerides were comparable between F/F and I-B/C KO mice (Fig. S6H), consistent with our observation that fatty acid absorption is unaffected by loss of Asters (Fig. S4A-D). These findings suggest that Aster deficiency reduces cholesterol content in chylomicrons, but does not alter the process of chylomicron assembly and release *per se*. In support of this conclusion, there were fewer ^14^C counts in the chylomicron fraction of plasma isolated 2 h after oral gavage of [^14^C]cholesterol, in I-B/C KO mice versus F/F mice (Fig. 3I). To show that this decrease in cholesterol was due to a reduction in CE in chylomicrons, compositional analysis of chylomicrons lipidome was performed in mice gavaged with d4-cholesterol and injected with Poloxamer-407. I-B/C KO mice displayed a reduction in d4-CE in isolated chylomicrons (Fig. 3J) without changes in d4- cholesterol compared to F/F mice (Fig. 3K). Unlabeled CE species were also lower, while no differences were observed in unlabeled cholesterol (Fig. S6I, J).

### Loss of Asters protects against diet-induced hypercholesterolemia

Next, we investigated whether intestinal Aster deficiency protects against diet-induced hypercholesterolemia. After 21 days of a HC diet, we observed a modest decrease in the weight of global B/C-KO mice (Fig. 4A), accompanied by reduced fasting plasma cholesterol levels (Fig. 4B). Liver cholesterol levels were also reduced in B/C-KO mice (Fig. 4C). We also found lower fasting plasma cholesterol levels in intestine-specific Aster-KO mice (Fig. 4E), with no changes in body weight (Fig. 4D). FPLC fractionation of the plasma revealed lower cholesterol levels in the VLDL/LDL and HDL fractions (Fig. 4F). Plasma triglyceride levels were similar in Aster knockout mice and controls (Fig. S7A and B). Gene expression (Fig. 4G and H) and western blot analyses (Fig. 4I) revealed an induction of the SREBP-2 pathway in global B/C-KO and I-B/C-KO mice. Thus, even with high levels of cholesterol in the diet, *de novo* cholesterol synthesis was activated in KO IECs. Lipidomic analyses in jejunal scrapings revealed reduced CE in I-B/C-KO enterocytes on the HC diet (Fig. 4J), without changes in other lipids species (Fig. S7C). We did not observe changes in body weight (Fig. S7D) or lipid levels in Aster single-KO mice after 21 days of HC diet (Fig. S7 E, F), nor did we observe an induction of SREBP-2 target genes (Fig. S7G).

**Fig. 4:**
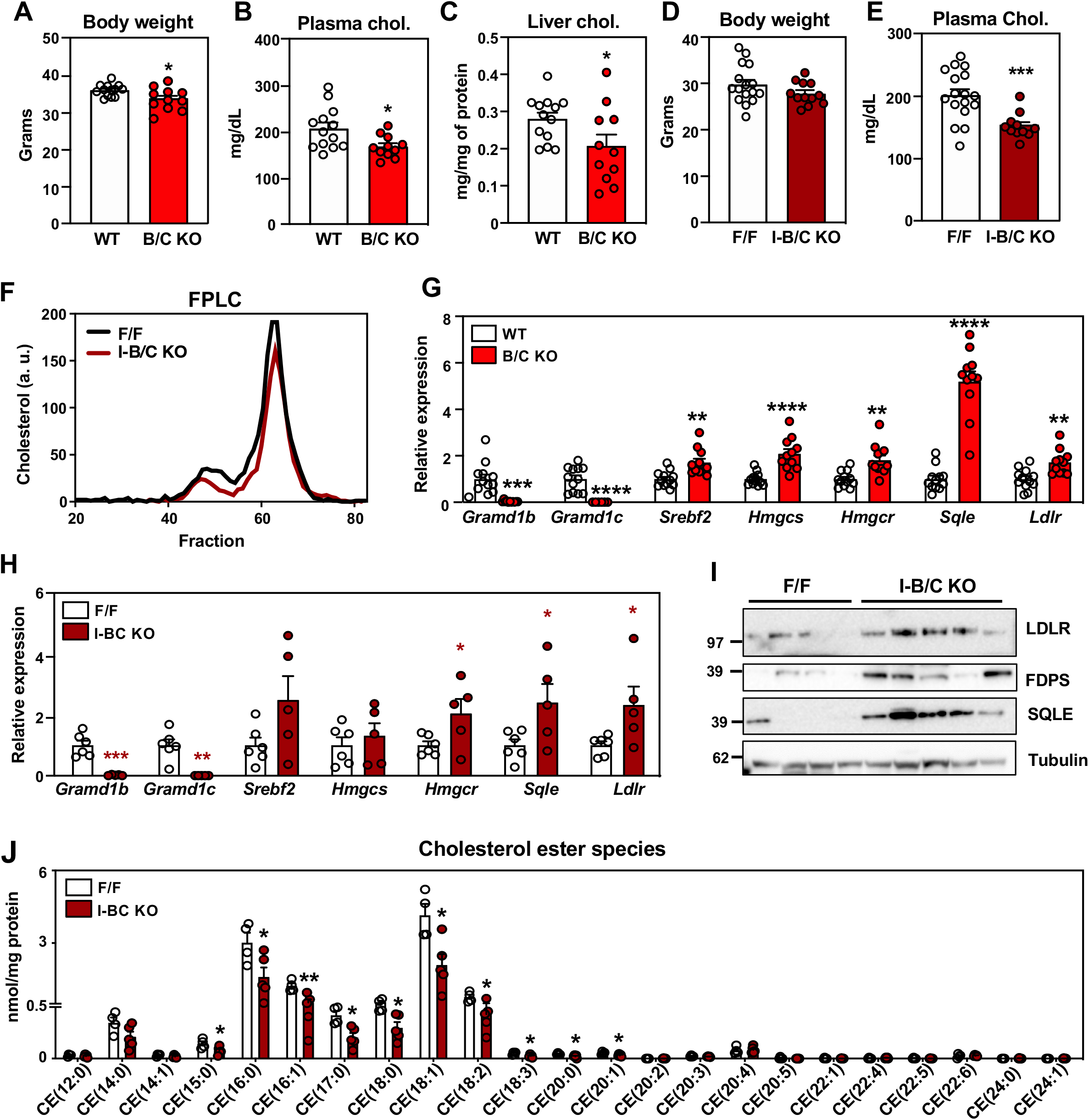
Deletion of intestinal Asters protects from diet induced hypercholesterolemia. (**A**) Body weight of WT (*n = 13*) and B/C KO (*n = 11*) male mice after 21 days of a western diet + 1.25% cholesterol (HC) diet; (**B**) Plasma cholesterol level after 4 h fasting in mice described in A; (**C**) Liver cholesterol in mice described in A; (**D**) Body weight of F/F (*n = 17*) and I-B/C KO (*n = 11*) male mice after 21 days of HC diet; (**E**) Plasma cholesterol level after 4 h of fasting in mice described in D; (**F**) FPLC of plasma from F/F and I-B/C KO mice fed for 21 days with HC diet and euthanized after overnight fasting followed by 2 h of refeeding with the HC diet (pool of 3–5/group); (**G**) Gene expression in distal jejunum scrapings from WT (*n = 12*) and B/C KO (*n = 11*) mice after 21 days of HC diet, euthanized after 4 h fasting; (**H**) Gene expression in distal jejunum scrapings from F/F (*n = 6*) and I-B/C KO (*n = 5*) male mice after 21 days of HC diet, euthanized after 4 h fasting; (**I**) Western blot analysis of duodenum scrapings of mice described in H (*n = 4/5*); (**J**) Lipidomic analysis of cholesterol esters in proximal jejunum scrapings from mice described in H (*n = 5*/group). Data are expressed as mean ± SEM. Statistical analysis: unpaired *t* test, *p < 0.05, **p < 0.01, ***p < 0.001, ****p < 0.0001.

### Ezetimibe binds to Aster-B and Aster-C

EZ reduces plasma cholesterol absorption by inhibiting the cholesterol transport function of NPC1L1. However, the mechanism by which EZ interferes with cholesterol delivery to the ER, and the effect of EZ on non-vesicular cholesterol transport, are incompletely understood (*7, 39*). Because both Aster deficiency and EZ reduce cholesterol absorption, we tested whether Asters might bind EZ. We previously showed that the ASTER domain binds cholesterol and other sterols (*27*). Using competition assays for 22-NBD-cholesterol binding and found that EZ bound to Aster-B and Aster-C (Fig. 5A) with moderate affinity, but exhibited minimal binding to Aster-A and StARD1 (Fig. S8A). Next, we solved the crystal structure, at 1.6 Å resolution, of the ASTER domain from Aster-C complexed to EZ. The overall structure of the Aster-C domain revealed a canonical curved seven-stranded beta-sheet that forms a cavity to accommodate EZ (Fig. 5B). The cavity is closed by a long carboxyl-terminal helix and two shorter helices. The structure highlighted additional volume within the binding cavity that accommodates a glycerol molecule, and density for part of a PEG4000 molecule from the cryo-protectant and crystallization buffer respectively (Fig. S8B, S8C). The electron density for the PEG is stronger at the end in proximity to the EZ ligand and becomes weaker at the other end, presumably due to disorder. Thus, the precise translational position of the PEG is uncertain. Modeling of ezetimibe- glucuronide, the active metabolite of EZ, in the pocket suggested potential capacity for binding, since the glucuronide group is oriented toward an opening of the pocket (Fig. S8D). Circular dichroism (CD) spectra of the Aster-A, -B, and -C domains revealed that Aster-B and -C were thermally stabilized by EZ binding, whereas Aster-A was not (Fig. S8E). Cholesterol and U18666A were included as positive controls (*27, 40*).

**Fig. 5:**
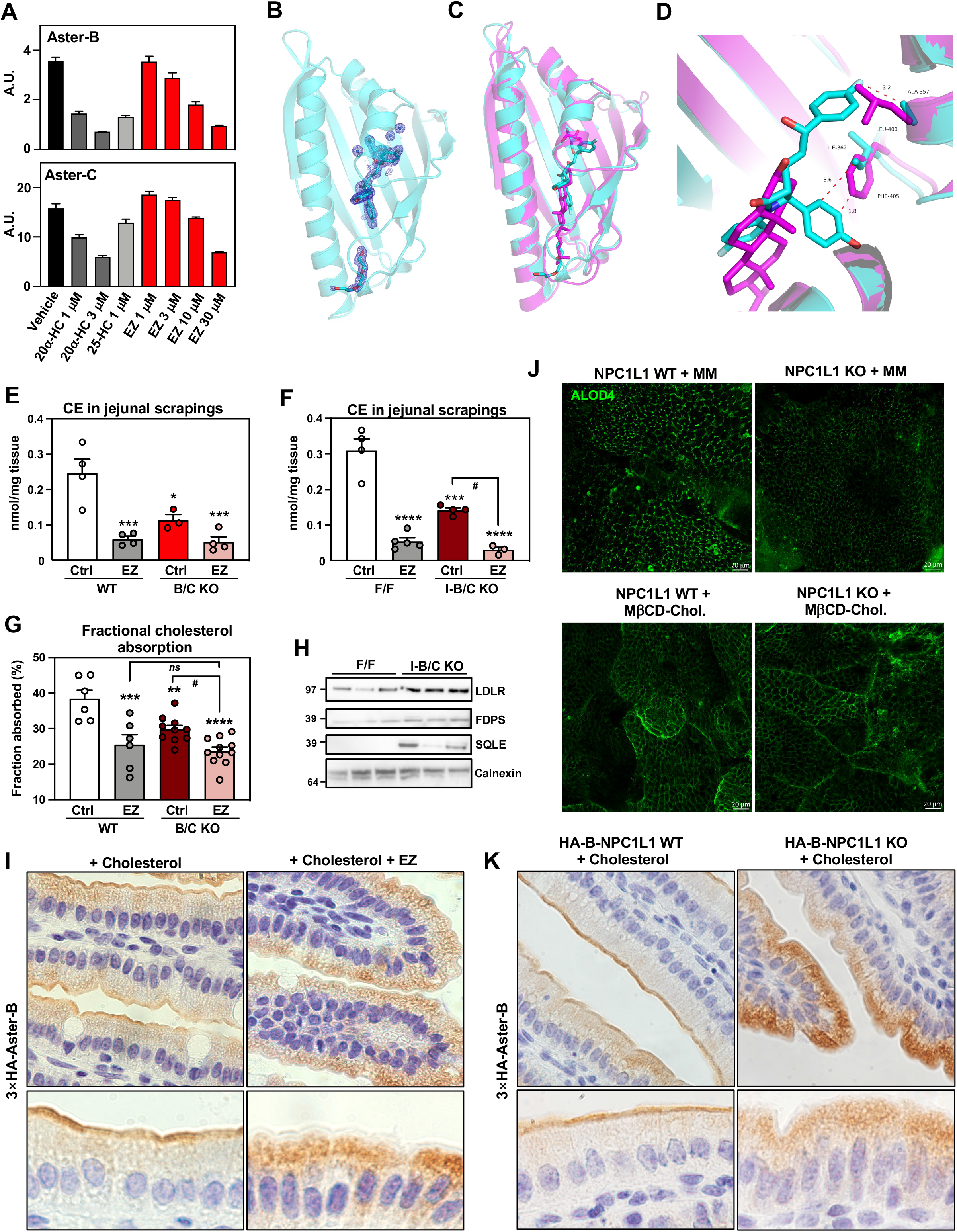
Asters bind ezetimibe and cooperate with NPC1L1. (**A**) Competition assays for 22-NBD-cholesterol binding to purified ASTER-B and ASTER-C domains incubated in presence of vehicle, 20α-HC, 25-HC, or EZ (1–30 mM). Data bars represent means ± SD; (**B**) Cartoon representation of the atomic structure, at 1.6 Å resolution, of the ASTER domain of Aster-C complexed to EZ. The EZ ligand in cyan sticks fits in a pocket created between the highly curved β-sheet, the second short helix and the carboxyl-terminal helix. The cavity also accommodates glycerol (cyan sticks), water molecules (blue spheres), and part of a PEG 4000 molecule (cyan sticks). 2Fo-Fc electron density map of EZ, glycerol, part of PEG 4000, and water molecules in the binding cavity shown as blue mesh contoured at 1.20; (**C**) Superposition of Aster-C:EZ (cyan) with the structure of Aster A:25-HC (pdb ID 6GQF). The overall architecture of the domain is very similar, but there are differences in the binding pocket and the opening loop; (**D**) Detail of the interaction between EZ and Aster-C involving residues ALA 357 and ILE 362, and superposition of the Aster-A:25-HC interaction. The superposition reveals how LEU 400 and PHE 405 from Aster A domain would clash with EZ, potentially preventing Aster A from binding EZ; (**E**) Cholesterol ester quantification by mass spectrometry in scrapings from proximal jejunum of WT and B/C KO in mice fed for 3 days with a control diet (ctrl) containing 0.08% cholesterol or a diet containing 0.08% cholesterol and 0.01% EZ and euthanized 2 h after refeeding (*n = 3-5*/group); (**F**) Cholesterol ester quantification by mass spectrometry in scrapings from proximal jejunum of F/F and I-B/C KO in mice fed for 3 days with a control diet (ctrl) containing 0.08% cholesterol or a diet containing 0.08% cholesterol and 0.01% EZ) and euthanized 2 h after refeeding (*n = 3-5*/group); (**G**) Fractional absorption of cholesterol measured by fecal dual isotope method in F/F and I-B/C KO mice fed for 3 days with a control diet (ctrl) containing 0.08% cholesterol or a diet containing 0.08% cholesterol and 0.01% EZ; (**H**) Western blot analysis of duodenum scrapings from F/F and I-B/C KO mice fed for 3 days with EZ diet (*n = 3*/group). This was run on the same membrane used for western blot analysis reported in Fig. S3G. Therefore, the loading control (Calnexin) is the same; (**I**) Immunohistochemistry of HA-Aster-B in small intestines from 3×HA-Aster-B mice after an oral administration of vehicle or EZ and a gastric gavage with cholesterol in corn oil; (**J**) ALOD4 imaging of murine enteroids from WT and NPC1L1 KO mice after loading with cholesterol in mixed micelles or with MbCD-cholesterol; (**K**) Immunohistochemistry of HA-Aster-B in small intestines from 3×HA-Aster-B crossed to NPC1L1 WT or NPC1L1 KO mice after a gastric gavage with cholesterol in corn oil. Data are expressed as mean ± SEM. 2-way ANOVA with Tukey’s as multiple comparisons test. Data are expressed as mean ± SEM. *p < 0.05, **p < 0.01, ***p < 0.001, ****p < 0.0001 vs WT Ctrl, ^#^p < 0.05 vs I-B/C KO Ctrl.

To investigate the molecular basis for selective binding of EZ to Aster-C, we compared the structures of Aster-C:EZ and Aster-A:25.-hydroxycholesterol (Fig. 5C). Many of the residues involved in ligand interactions are conserved in Aster-A and Aster-C (Fig. S8F). However, leucine 400 and phenylalanine 405 in the ASTER domain of Aster-A appeared to represent a steric hindrance to EZ binding (Fig. 5D), while alanine 357 and isoleucine 362 in the ASTER domain of Aster-C left ample space for EZ binding (Fig. 5D). The importance of the specific amino acids on EZ binding was interrogated by mutagenesis and evaluating thermal stability by CD. The two residues that appeared to hinder EZ binding in Aster-A were changed to the residues found in Aster-C (L400A_F405I). Conversely, the residues that appeared to be important for accommodating EZ binding in Aster-C were changed to those in Aster-A (A357L_I362F) which potentially prevent binding. The Aster-A mutant L400A_F405I, as well as the single mutant L400A, reduced Aster-A thermal stability (Fig. S8E), but the stability was increased by EZ. The thermal stability upon EZ binding was unchanged in the F405I mutant (Fig. S8G), suggesting that L400A is crucial for binding selectivity. These results were corroborated by the Aster-C mutants, where A357L_I362F and A357L alone increased thermal stability of the protein, while I362F had no effect. The thermal stability of the A357L Aster-C mutant did not change with EZ binding, implying a weak interaction between EZ and that mutant (Fig. S8G).

### Intestinal Asters cooperate with NPC1L1 in intestinal cholesterol absorption

The ability of Aster-B and -C to bind EZ suggested that they may act in concert with NPC1L1 to promote cholesterol absorption and efficient packaging into chylomicrons. To test this hypothesis, we fed WT and Aster-deficient mice a moderate-cholesterol control diet (Ctrl) or a moderate-cholesterol diet containing 0.01% EZ diet for 3 days (Fig. S9A). As expected, EZ decreased CE content and fractional cholesterol absorption by the fecal dual isotope method in WT mice (Fig. 5E-G). Aster-B/C-KO and I-B/C-KO mice showed decreased CE content (Fig. 5E, F) and fractional cholesterol absorption (Fig. 5G); EZ treatment led to a further reduction. On the other hand, Aster deficiency did not provide an additional decrease in CE or cholesterol absorption in EZ-treated mice (Fig. 5E-G). However, gene-expression analyses revealed stronger activation of the SREBP-2 pathway in Aster-B/C-KO mice on the EZ diet compared to either Aster-B/C KO mice or EZ-treated mice (Fig. S9B). Thus, the combination of Aster deficiency and EZ treatment resulted in an additive or synergistic induction of *de novo* cholesterol synthesis genes. Western blot analyses revealed that protein levels of SREBP-2 targets were induced in a similar manner (Fig 5H).

We showed above that cholesterol loading triggers Aster-B movement to the PM in enteroids (Fig. 1C). A prior study showed that EZ impairs cholesterol uptake by blocking the channel within NPC1L1 that is required for cholesterol deposition into the PM (*7*). Therefore, we theorized that EZ treatment would attenuate Aster translocation to the PM by preventing the expansion of accessible cholesterol pool at the PM. To test this idea, 3×HA-Aster-B mice were fed Ctrl diet or EZ diet for 3 days and gavaged with vehicle or EZ 30 minutes prior to a second gavage of cholesterol in corn oil. Small intestines were harvested 1 h later.

Immunohistochemical staining for HA-Aster-B revealed that EZ completely prevented the recruitment of Aster-B to the brush border of enterocytes after cholesterol gavage (Fig. 5I). Next, we everted enteroids such that the apical PM was facing out (*41*) and administered mixed micelles (MM) containing cholesterol in the presence of vehicle or EZ. Immunofluorescence microscopy confirmed that recruitment of HA-Aster-B to the apical PM of enteroids was induced by cholesterol loading in MM but reduced by EZ (Fig. S9C). In enteroids derived from NPC1L1-KO mice loaded with cholesterol in mixed micelles, we detected decreased ALOD4 staining, indicating that NPC1L1 is required to saturate the PM with cholesterol (Fig. 5J, upper panels). Interestingly, when enteroids were loaded with MβCD cholesterol, which directly deposits cholesterol into the PM and presumably bypasses NPC1L1, ALOD4 binding was not different between NPC1L1KO and WT enteroids (Fig. 5J, bottom panels).

We then crossed 3×HA-Aster-B mice to NPC1L1 WT (HA-B-NPC1L1 WT) or NPC1L1 KO (HA-B-NPC1L1 KO). In enteroids derived from these mice, NPC1L1 deletion reduced 3×HA-Aster-B recruitment to the PM after cholesterol loading with MM, while NPC1L1 was dispensable for Aster-B movement to the PM in enteroids loaded with MβCD cholesterol (Fig. S9D). IHC studies with HA-B-NPC1L1 WT and HA-B-NPC1L1 KO mice confirmed that genetic ablation of NPC1L1 abolished Aster-B movement to the brush border in response to cholesterol loading, mirroring the effects of EZ (Fig. S9E and Fig. 5K). These results substantiate the hypothesis that EZ prevents Aster translocation to the brush border via NPC1L1 blockade. EZ thus prevents the cholesterol saturation of the apical membrane by inhibiting NPC1L1 rather than via inhibiting of Aster-B.

Next, we generated a cell line of McA-RH7777 CRL-1601 hepatocytes stably expressing NPC1L1-EGFP and HA-Aster-B fusion proteins. Following cholesterol depletion by incubation in LPDS, NPC1L1 partially localized to PM, while Aster-B was confined to the ER (Fig. S9F). MβCD-cholesterol loading caused internalization of NPC1L1-EGF and, concomitantly, the movement of HA-Aster-B to ER-PM contact sites. This contrasting behavior of NPC1L1 and Aster in response to cholesterol loading argues against a direct physical interaction between the two proteins in cellular cholesterol import.

### Pharmacological Aster inhibition reduces intestinal cholesterol uptake

Previous work from our laboratory identified a small molecule, AI-3d, that potently inhibits Aster-A, -B, and -C (*40*). We tested this molecule in enteroids and *in vivo* to determine its ability to mimic the effects of Aster deficiency. First, we pretreated WT and B/C-KO enteroids with AI-3d and assessed accessible cholesterol at the PM with the ALOD4 probe.

Treatment of WT enteroids with AI-3d and MβCD cholesterol led to an accumulation of ALOD4 compared to untreated WT enteroids; while AI-3d had no additional effect on ALOD4 binding in B/C-KO enteroids (Fig. 6A). AI-3d displayed a similar effect on human jejunal enteroids (Fig. 6B), indicating that AI-3d inhibits both human and mouse Asters. We also compared the effect of AI-3d on WT and NPC1L1 KO enteroids. When enteroids were loaded with cholesterol in MM, AI-3d treatment increased ALOD4 binding only in NPC1L1 WT cells (Fig. S10A). However, when enteroids were loaded with MβCD-cholesterol, AI-3d treatment increased ALOD4 binding similarly in WT and NPC1L1 KO cells (Fig. S10B).

**Fig. 6:**
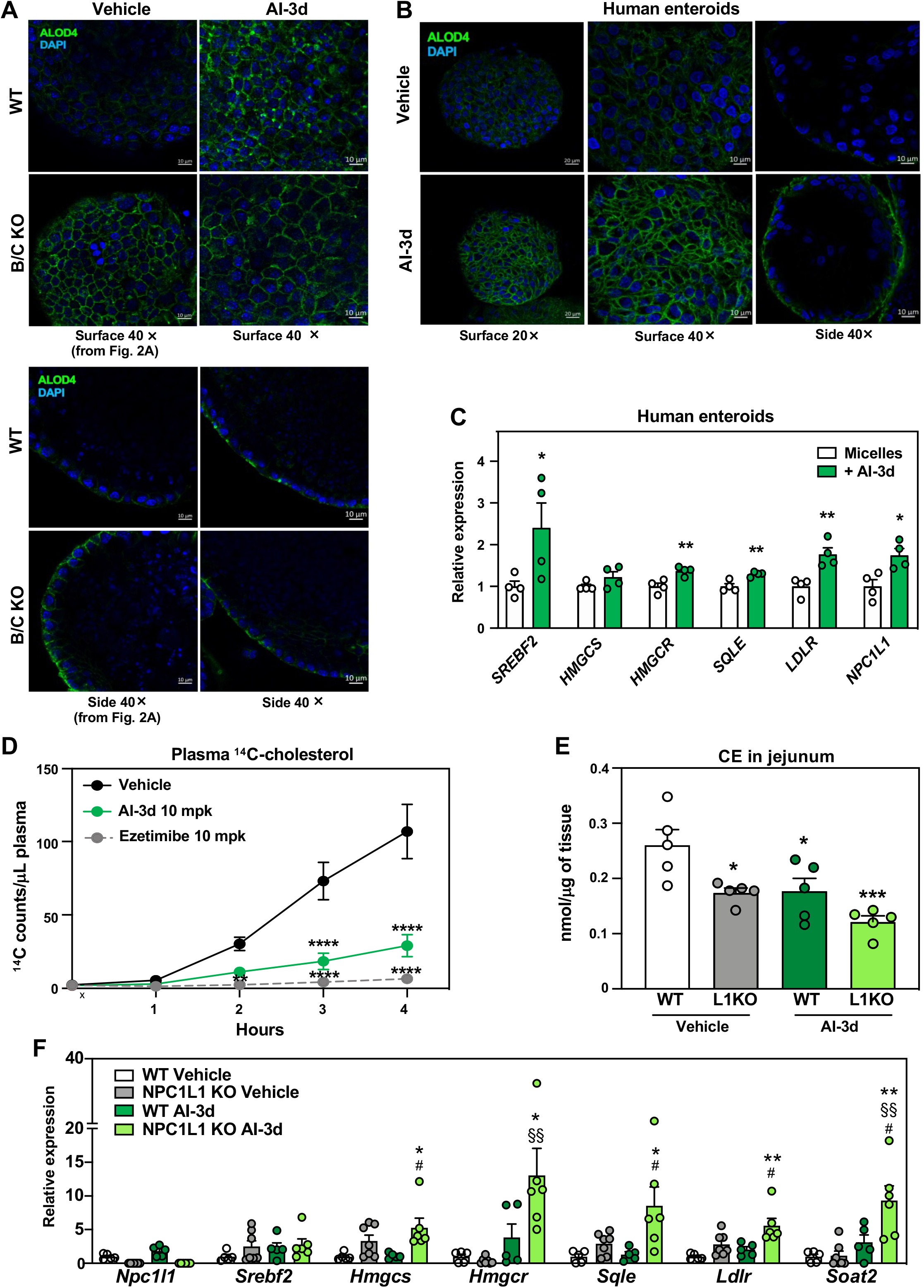
Pharmacological inhibition of Asters reduces intestinal cholesterol uptake. (**A**) ALOD4 staining (green) of PM accessible cholesterol in enteroids from WT and B/C KO treated with vehicle or Aster inhibitor AI-3d. Results presented in this panel and in Fig.2A came from one experiment, where WT vehicle-treated enteroids served as control for both B/C KO vehicle-treated enteroids and for AI-3d vehicle-treated enteroids; (**B**) ALOD4 staining (green) of PM accessible cholesterol in human enteroids treated with vehicle or AI-3d; (**C**) Gene expression of SREBP-2 and target genes in human enteroids differentiated on transwell and loaded with cholesterol in mixed micelles in the presence of vehicle or AI-3d; (**D**) Kinetics of radioactivity in plasma of female mice that were given an oral administration of vehicle (*n = 6*), AI-3d (*n = 5*), or EZ (*n = 6*) in corn oil. After 1 h, the mice were given a gastric gavage of olive oil containing [^14^C]cholesterol; (**E**) Cholesterol ester quantification by mass spectrometry in scrapings from proximal jejunum of NPC1L1 WT and NPC1L1 KO mice fed for 3 days with a control diet (ctrl) containing 0.08% cholesterol and treated with 3 doses of vehicle (left) or 10 mg/kg AI-3d (right) and euthanized 2 h after refeeding (*n = 5*/group); (**F**) Gene expression analysis of distal jejunum scrapings from mice described in E. Statistical analysis for panel C, unpaired *t* test; for panels, D, E, F, 2-way ANOVA with Tukey’s as multiple comparisons test. Data are expressed as mean ± SEM. For panel C, *p < 0.05, **p < 0.01 vs micelles. For panel D **p < 0.01, ****p < 0.0001 vs Vehicle. For panels D, E *p < 0.05, **p < 0.01 vs WT Vehicle, **^#^**p < 0.05 vs WT AI-3d, **^§§^**p < 0.01 vs NPC1L1 KO Vehicle.

Next, we assessed the impact of AI-3d on cholesterol transport to the ER. Consistent with our results in Aster-B/C KO and I-B/C KO mice, AI-3d led to an increase in SREBP-2 target gene expression in human enteroids (Fig. 6C) and in differentiated Caco-2 cells (Fig. S10C). These findings indicate the depletion of ER cholesterol in the presence of AI-3d.

We further tested the effect of AI-3d on cholesterol absorption *in vivo*. C57BL/6J mice were pretreated with AI-3d or EZ for 1 h, gavaged with [^14^C]cholesterol, and given an intraperitoneal injection of Poloxamer-407. EZ reduced the appearance of [^14^C]cholesterol in the plasma as expected, but interestingly AI-3d had a similar effect (Fig. 6D). This finding suggests that pharmacologic inhibition of Aster proteins reduces dietary cholesterol absorption.

Above we showed that the combination of EZ treatment and Aster deficiency led to a greater decrease in cholesterol absorption than Aster deficiency alone. Next, we evaluated the impact of Aster inhibition on WT and NPC1L1 KO mice fed a moderate-cholesterol diet. Mice received 3 doses of vehicle (corn oil) or 10 mg/kg AI-3d (Fig. S10D). AI-3d did not affect body weight or intestinal length (Fig. S10 E-G), supporting the absence of toxicity of this regimen.

Lipidomic analysis revealed reduction of CE stores in jejunal scrapings from NPC1L1 KO compared to WT mice. Aster inhibition by AI-3d lowered the levels of jejunal CE similarly in WT and NPC1L1 KO, indicating that AI-3d does not act through NPC1L1 (Fig. 6E). Gene expression analysis showed a further induction of SREBP2 targets in NPC1L-KO mice treated with AI-3d (Fig. 6F), an effect reminiscent of the transcriptomic effects induced by concomitant genetic deletion of Asters and pharmacologic inhibition of NPC1L1 by EZ (Fig. S9B).

Finally, we tested the effect of AI-3d on NPC1L1 recycling using a McA-RH7777 CRL- 1601 stably expressing NPC1L1-EGFP fusion protein. Consistent with published data (*17*), we observed that NPC1L1-EGFP was present within the endocytic recycling compartment (ERC) when both vehicle-treated and AI-3d treated cells were cultured in full serum (Fig. S10H, upper panels). In response to cholesterol depletion by MβCD the signal in both cell types moved to the PM (Fig. S10H, middle panels). Conversely, MβCD-cholesterol loading led to the relocation of NPC1L1-EGFP to the ERC (Fig. S10H, lower panels). These results indicate that AI-3d treatment does not interfere with the process of NPC1L1 recycling, further supporting the conclusion that AI-3b inhibits cholesterol absorption by targeting Asters.

## Discussion

The involvement of NPC1L1 in facilitating the entry of cholesterol into enterocytes and the role of ACAT2 in cholesterol esterification are well documented, and both proteins are recognized to be key players in the physiological process of intestinal cholesterol absorption (*3, 12*). However, how cholesterol that enters the cell via NPC1L1 reaches the ER for esterification and regulation of cholesterol synthesis has been a long-standing mystery. Here we solve that mystery by showing that Aster-B and -C link NPC1L1 to ACAT2 by facilitating cholesterol transport to the ER following NPC1L1 uptake in enterocytes.

Our current studies demonstrate that combined deletion of Aster-B and -C impairs the movement of dietary cholesterol from apical PM to ER, as evidenced by expansion of the pool of accessible cholesterol at PM and by reduced CE formation in the ER. Physiologically, the impaired transport of cholesterol from the PM to the ER reduces cellular cholesterol stores and impairs the incorporation of CE into chylomicrons. At a molecular level, the reduced delivery of PM cholesterol to the ER in the setting of Aster deficiency activates the SREBP-2 pathway, leading to increased transcription of the genes driving cholesterol synthesis. The activation of *de novo* cholesterol synthesis likely compensates for the excessive drop in CE stores and allows chylomicron production to proceed. Interestingly, the degree of activation of the SREBP-2 pathway in Aster-deficient mice was even greater when cholesterol entry into the apical PM was blocked by EZ. Similarly, genetic ablation of NPC1L1 combined with pharmacological inhibition of Asters led to further induction of SREBP-2 target genes. These observations imply a particularly important function for Asters in cholesterol delivery to the ER, and suggest that they may accept cholesterol from sources other than the apical PM, such as from the basolateral PM where NPC1L1 is absent.

We found that EZ binds to Aster-B and -C with moderate affinity, raising the possibility that EZ-mediated inhibition of Aster function could contribute to EZ’s capacity to inhibit cholesterol absorption. However, EZ binding to Aster is not sufficient to block non-vesicular cholesterol transport *in vivo*, as Aster deletion further reduces ER cholesterol delivery in the presence of EZ. These findings imply that the binding of EZ to Aster proteins does not inactivate the function of the Aster proteins. Our results suggest that cholesterol homeostasis in enterocytes requires both NPC1L1-mediated cholesterol deposition into the PM of enterocytes and subsequent Aster-mediated cholesterol transport to the ER. Since NPC1L1 is the gatekeeper that controls the first step of cholesterol uptake, when its function is blocked by EZ cholesterol absorption is reduced. In this context, the deletion of Asters does not further decrease absorption, because the amount of apical PM cholesterol available for the transfer to ER is limited.

The recruitment of Asters to the PM depends on the cholesterol and phosphatidylserine content in the inner leaflet of the plasma membrane (*27, 28, 35*). Our results support the notion that Aster-mediated non-vesicular cholesterol transport relies on NPC1L1 to deposit cholesterol into the apical PM, rather than on a direct physical interaction with NPC1L1. In support of this concept, micelle-derived cholesterol cannot contribute to the expansion of accessible pool and Aster proteins do not translocate to the enterocyte PM when NPC1L1 is knocked out or is inhibited by EZ. However, when NPC1L1 is bypassed by delivering cholesterol directly to the PM with MβCD, Aster is recruited normally. These observations are in line with our previous studies of Asters in cells that do not express NPC1L1, where saturation of membrane with cholesterol is mediated by other players, such as SR-BI. In enterocytes, where NPC1L1- mediated deposition of cholesterol is the primary mechanism to expand the accessible cholesterol pool, Asters co-operate with NPC1L1 to allow a rapid delivery of cholesterol to ER. Our studies do not exclude the possibility that NPC1L1 internalizes cholesterol and that Aster proteins then retrieve that cholesterol, directly or indirectly, from NPC1L1-rich endosomes.

Our findings indicate that ASTER-mediated cholesterol transport could be a target for treating diet-induced hypercholesterolemia. First, we demonstrated that Aster-B/C knockout mice have low plasma cholesterol levels when fed a western diet enriched in cholesterol. Second, we showed that the Aster pathway can be targeted pharmacologically to reduce cholesterol absorption. The Aster inhibitor AI-3d, which had been shown previously to inhibit Aster-mediated non-vesicular transport *in vitro*, reduced cholesterol transport to the ER in both mouse and human enteroids. AI-3d also expanded the pool of accessible cholesterol at the plasma membrane of intestinal enteroids. Finally, treatment of mice with AI-3d inhibited cholesterol absorption at levels comparable to those observed with EZ. These findings identify the Aster pathway as a potentially attractive pathway for limiting intestinal cholesterol absorption and reducing plasma cholesterol levels.

## Acknowledgments

We thank Loren Fong, Paul Kim and Elmira Tokhtaeva for the technical assistance. We also thank all members of the Tontonoz, Tarling-Vallim, Edwards, Villanueva, Young and Bensinger labs at UCLA for useful advice and discussions and for sharing reagents and resources. We thank G. Su and the UCLA Lipidomics core for the lipidomics analysis, and Y. Li and the UCLA Translational Pathology Core Laboratory for tissue histology and immunohistochemistry. We are in debt with Bao Liang Song at Wuhan University for sharing NPC1L1-EGFP construct and NPC1L1 KO mice. We thank Tamim Darwish for supporting the production of [2H]cholesterol. The National Deuteration Facility in Australia is partly funded by The National Collaborative Research Infrastructure Strategy (NCRIS), an Australian Government initiative.

## Author contributions

Conceptualization: AF, EW, PT

Methodology: AF, EW, XX, JPK, SDL, AN, TW, LF, MGM, RAR, LC, AB, KW

Investigation: AF, EW, XX, JPK, BRA, JJM, TW, KC, JS, MJT, LB, AN, YK, YG, PM, WS

Visualization: AF, EW, PT

Funding acquisition: SGY, PT

Project administration: PT

Supervision: MEJ, HJ, JWRS, SGY, PT

Writing – original draft: AF, EW, PT

Writing – review & editing: AF, EW, MEJ, HJ, JWRS, SGY, PT

## Data and materials availability

Sequencing data have been deposited to GEO (GSE206780). The crystal structure has been submitted to the PDB database, PDB ID code 8AXW.

## Supplementary material

### Summary

Materials and Methods

Figs. S1 to S11

Tables S1 to S2

References (*40–58*)

