## Supplemental material for "Aster-dependent non-vesicular transport facilitates dietary cholesterol uptake"

Science
AAAS

### Supplementary Materials for

#### Aster-dependent non-vesicular transport facilitates dietary cholesterol uptake

Alessandra Ferrari†, Emily Whang†, Xu Xiao, John P. Kennelly, Beatriz Romartinez-Alonso, Julia Mack, Thomas Weston, Kai Chen, Youngjae Kim, Marcus J. Tol, Lara Bideyan, Alexander Nguyen, Yajing Gao, Liujuan Cui, Alexander H. Bedard, Jaspreet Sandhu, Stephen D. Lee, Louise Fairall, Kevin J. Williams, Wenxin Song, Priscilla Munguia, Robert A. Russell, Martin G. Martin, Michael E. Jung, Haibo Jiang, John W.R. Schwabe, Stephen G. Young, Peter Tontonoz*

**This PDF file includes:**

Materials and Methods
Figs. S1 to S8
Table S1 to S2

**Other Supplementary Materials for this manuscript include the following:**

References (*40*-*58*)

#### Supplemental Figure Legends

**Fig. S1. Expression and regulation of Aster variants in small intestine**

(**A**) RNA sequencing tracks of *Gramd1a*, *Gramd1b*, and *Gramd1c* from proximal jejunum of WT mouse; (**B**) Quantification of *Gramd1a*, *Gramd1b*, and *Gramd1c* transcripts in isolated intestinal epithelial cells (IECs) ($n = 5$); (**C**) Gene expression of differentiation markers and *Gramd1a*, *Gramd1b*, and *Gramd1c* in Caco-2 cells that had been differentiated on transwells; (**D**) Gene expression of LXR targets and *Gramd1a*, *Gramd1b*, and *Gramd1c* in jejunum scrapings from mice treated with vehicle ($n = 6$) or the LXR ligand GW3965 ($n = 4$) for 8 h; (**E**) Generation of 3×HA-Aster-B mice by CRISPR/Cas9 based strategy; (**F**) Detection of 3×HA-Aster-B in small intestine by western blot; (**G**) Immunohistochemistry of HA-Aster-B in small intestines from 3×HA-Aster-B mice, showing higher expression in the villi. Data are expressed as mean ± SEM. Statistical analysis: unpaired *t* test, $^*p < 0.05$, $^{**}p < 0.01$, $^{***}p < 0.001$.

**Fig. S2. Preserved cholesterol absorption in global Aster-B and Aster-C single KO mice.**

(**A**) Generation of Aster-B global KO (BKO) mice; (**B**) Generation of Aster-C global KO (CKO) mice; (**C**), (**D**), (**E**) Deletion of Aster-B and Aster-C in single and double KO confirmed by quantitative PCR ($n = 3–5$/group); (**F**) Hematoxylin and eosin–staining of proximal jejunum of WT and Aster-B/C global knockout (B/C KO) mice (**G**) Body weight and intestinal length of BKO, CKO, and B/C KO mice compared to littermates WT ($n = 4–9$/group); (**H**) Distribution of radioactivity in intestinal segments of female WT and BKO mice after an oral challenge of olive oil containing [$^{14}$C]cholesterol for 2 h ($n = 4–5$/group); (**I**) Radioactivity in liver of mice described in I; (**J**) Radioactivity in plasma of mice described in I; (**K**) Distribution of radioactivity in intestinal segments of female WT and CKO mice after an oral challenge of olive oil containing [$^{14}$C]cholesterol for 2 h ($n = 8–9$/group); (**L**) Radioactivity in liver of mice described in L; (**M**) Radioactivity in plasma of mice described in L. Data are expressed as mean ± SEM. Statistical analysis: unpaired *t* test, $^*p < 0.05$, $^{**}p < 0.01$, $^{***}p < 0.001$.

**Fig. S3. Preserved cholesterol absorption in inducible Aster-B and Aster-C single-KO mice.**

(**A**) Generation of intestinal tamoxifen inducible Aster-B conditional KO (I-BKO) mice; (**B**) Generation of intestinal tamoxifen inducible Aster-C conditional KO (I-CKO) mice; (**C**), (**D**), (**E**) Deletion of Aster-B and Aster-C in single and double conditional KO mice confirmed by quantitative PCR ($n = 3–5$/group); (**F**) Western blot analysis confirmed deletion of Aster-B in I-B/C KO mice (n = 3/group); (**G**) Immunohistochemistry of Olfm4 in jejunum of F/F and I-B/C KO; (**H**) Body weight and intestinal length of I-BKO, I-CKO, and I-B/C KO mice compared to F/F littermates ($n = 6–10$/group); (**I**) Distribution of [$^{14}$C]cholesterol radioactivity in intestinal segments of male F/F and I-BKO mice after an oral challenge of olive oil containing [$^{14}$C]cholesterol for 2 h ($n = 6–8$/group); (**J**) Radioactivity in liver of mice described in I; (**K**) Radioactivity in plasma of mice described in I; (**L**) Distribution of radioactivity in intestinal segments of female F/F and I-CKO mice after an oral challenge of olive oil containing [$^{14}$C]cholesterol for 2 h ($n = 7–10$/group); (**M**) Radioactivity in liver of mice described in K; (**N**) Radioactivity in plasma of mice described in L. Data are expressed as mean ± SEM. Statistical analysis: unpaired *t* test; $^*p < 0.05$, $^{**}p < 0.01$, $^{***}p < 0.001$.

**Fig. S4. Fatty acid absorption is intact in intestine-inducible Aster-B/C KO mice**

(**A**) Distribution of radioactivity in intestinal segments of female F/F and I-B/C KO mice after an oral challenge of olive oil containing [$^{14}$C]triolein for 2 h ($n = 6$/group); (**B**) Radioactivity in liver of mice described in A; (**C**) Radioactivity in liver of mice described in A; Cumulative radioactivity in intestinal segments of mice described in (A); (**D**) Glucose uptake, measured as plasma glucose appearance in female F/F and I-B/C KO mice after oral gavage of 2 mg/kg glucose. Data are expressed as mean ± SEM. Statistical analysis: for panels A, D, E, 2-way ANOVA with Tukey's as multiple comparisons test; for panels B, C, unpaired *t* test; *p < 0.05, **p < 0.01, ***p < 0.001.

**Fig. S5. Intestinal deletion of Asters reduces accumulation of dietary derived cholesterol in distal duodenum.**
(**A**) Backscattered electron image and NanoSIMS images of the distal duodenum from WT and B/C KO mice after having been given, by gastric gavage, a mixture of [$^{2}$H]cholesterol and [$^{13}$C]mixed fatty acids; (**B**) Quantification of $^{2}H^{-}$ secondary ions from the intestinal villus, relative to $^{1}H^{-}$, $^{13}C^{-}$, and $^{16}O^{-}$ secondary ions. Data are expressed as mean ± SEM. Statistical analysis: unpaired *t* test; *p < 0.05, ***p < 0.001, ****p < 0.0001.

**Fig. S6. Characterization of phenotypical effect of Aster deletion on intestinal cholesterol metabolism**
(**A**) Lipidomic analysis in proximal jejunum of WT (*n = 3*) and B/C KO (*n = 4*) mice after refeeding for 2 h with chow diet; (**B**) Quantification of $^{14}$C in proximal jejunum scrapings of WT (*n = 5*) and B/C KO (*n = 4*) 2 h after oral gavage with [$^{14}$C]cholesterol; (**C**) Gene-expression analysis in distal jejunum of WT (n = 4) and B/C KO mice (*n = 4*) 2 h after refeeding with western diet + 1.25 % cholesterol; (**D**) Gene-expression analysis in distal jejunum of WT (*n = 9*) and BKO (*n = 7*) mice after 4 h of fasting; (**E**) Gene-expression analysis in distal jejunum of WT (*n = 10*) and CKO (*n = 7*) mice after 4 h of fasting; (**F**) Gene-expression analysis in distal jejunum of F/F (*n = 3*) and I-BKO (*n = 3*) after 4h fasting; (**G**) Gene-expression analysis in distal jejunum of WT (*n = 4*) and CKO (*n = 7*) mice after 4 h of fasting; (**H**) Plasma triglycerides levels B/C-KO mice after refeeding a HC diet for 2 h; (**I**), (**J**) Unlabeled cholesterol esters and unlabeled cholesterol in chylomicrons isolated from plasma of F/F (n = 4) and I-B/C KO (*n* = 4) 3.5 h after treatment with Poloxamer-407 and oral gavage of cholesterol-d4. Data are expressed as mean ± SEM. Statistical analysis: unpaired *t* test; **p < 0.01.

**Fig. S7. Global BKO and CKO are not protected from diet induced hypercholesterolemia**
(**A**) Fasting plasma triglycerides in WT (*n = 13*) and B/C KO (*n = 11*) male mice after 21 days of the HC diet; (**B**) Fasting plasma triglycerides in F/F (*n = 17*) and I-B/C KO (*n = 11*) male mice after 21 days on the HC diet; (**C**) Lipidomic analysis from proximal jejunum of F/F (*n = 4*) and I-B/C KO (*n = 5*) male mice fed HC diet for 21 days and euthanized after 4 h of fasting; (**D**) Body weight of WT (*n = 13*), BKO (*n = 7*), and CKO (*n = 10*) mice after 21 days of the HC diet; (**E**) Fasting plasma cholesterol levels in mice described in (D); (**F**) Fasting plasma triglyceride levels for mice described in (D); (**G**) Gene expression in distal duodenum scrapings of mice described in (C). Data are expressed as mean ± SEM. Statistical analysis: unpaired *t* test; *p < 0.05, **p < 0.01.

**Fig. S8. Selective binding of ASTER-B and -C to ezetimibe**
(**A**) Competition assays for 22-NBD-cholesterol binding to purified ASTER-A and STAR domains incubated in presence of vehicle, 20α-HC, 25-HC, EZ (1–30 mM). Bars represent means ± SD; (**B**) 2Fo-Fc electron density map (contoured at 1.2σ) of Aster-C:EZ showing the ligand EZ along with glycerol and part of PEG 4000 (in sticks), residues surrounding them (in

grey sticks) and water molecules (blue spheres) displayed as blue mesh;; (**C**) Simulated annealing composite omit map (contoured at 1.2σ of Aster-C:EZ showing EZ, glycerol and part of PEG 4000 (in sticks), residues surrounding them (in grey sticks) and water molecules (blue spheres) displayed as blue mesh; (**D**) Modeling of ezetimibe-glucuronide in the pocket showing that glucuronide group is oriented toward an opening of the pocket; (**E**) Graphic representation of the ellipticity vs. temperature circular dichroism findings of Aster-A, -B and -C in the absence of ligand (blue), or in the presence of several ligands (magenta, 25-HC cholesterol; cyan, EZ; purple, U18666A); (**F**) Alignment of Aster-A, -B, and -C domains, highlighting the residues that interact with 25-HC in Aster-A (magenta) and with EZ in Aster-C (cyan). Red arrows point to the main differences in residues involved in ligand interactions in the three proteins; (**G**) Thermal stability of wild-type and mutant Aster-A and -C. The graph shows the melting temperature of the different proteins in the presence or absence of ligands with a statistical comparison between the apo wild-type and the mutants, and between the apo and the ligands within the same protein. Data are expressed as mean ± SEM. Statistical analysis: One-way ANOVA, with Dunnet as post hoc test; $*p < 0.05$, $**p < 0.01$, $***p < 0.001$, $****p < 0.0001$.

**Fig. S9. Mechanism of cooperation of NPC1L1 and Asters to uptake cholesterol and deliver to ER**

(**A**) Experimental details for the exposure of WT, B/C KO, F/F, and I-B/C KO to a control diet or to a diet containing 0.01% EZ; (**B**) Heatmap showing expression of ABC transporters, *Npc1l1*, *Soat2,* and genes involved in the *de novo* synthesis of cholesterol in distal jejunum scrapings from WT and B/C KO mice2 hours post refeeding (*n = 3-4*/group); (**C**) Confocal immunofluorescence micrographs of the apical marker ZO-1 and HA-Aster-B in enteroids isolated from 3×HA-Aster-B ileum and grown apical out showing recruitment of HA-Aster-B to the apical membrane without cholesterol loading, with cholesterol loading in mixed micelles in presence of vehicle or EZ; (**D**) Immunofluorescence microscopy of HA-Aster-B in intestinal organoids from HA-B-NPC1L1 WT or HA-B-NPC1L1 KO mice during loading with MbCD-cholesterol or cholesterol in mixed micelles; (**E**) Immunohistochemistry of HA-Aster-B in small intestines from 3×HA-Aster-B crossed to NPC1L1 WT or NPC1L1 KO mice after a gastric gavage with corn oil; (**F**) Confocal imaging and immunofluorescence in a stable cell line expressing NPC1L1-EGFP and HA-Aster-B fusion proteins, during cholesterol depletion by LPDS or MβCD-cholesterol loading.

**Fig. S10. Pharmacological modulation of Aster function by small molecule inhibitor AI-3d**

(**A**) ALOD4 imaging of murine enteroids from WT and NPC1L1 KO mice, treated with vehicle or AI-3d, after loading with cholesterol in mixed micelles; (**B**) ALOD4 imaging of murine enteroids from WT and NPC1L1 KO mice, treated with vehicle or AI-3d, after loading or with MβCD-cholesterol; (**C**) Gene expression analysis of SREBP2 and target genes in Caco-2 cells differentiated on transwell and loaded with cholesterol in mixed micelles in the presence of vehicle or AI-3d; (**D**) Experimental details for the exposure of WT and NPC1L1 KO to a low cholesterol diet orally gavaged with vehicle or 3 doses of 10mg/kg in corn oil; (**E**) Body weight of mice described in C; (**E**), (**F**) Intestinal length for mice described in C; **H**) McA-RH7777 CRL-1601 hepatocytes stably expressing NPC1L1-EGFP fusion protein, in cells cultured in full serum, after cholesterol depletion with MβCD, or cholesterol depletion with MβCD followed by loading with MβCD-cholesterol for 2 h; Data are expressed as mean ± SEM. Statistical analysis:

for panel C, unpaired *t* test; for panels E, F, G, 2-way ANOVA with Tukey’s as multiple comparisons test *$p < 0.05$, **$p < 0.01$.

#### Materials and Methods

Mice

Aster-B global knock out mice were described before (*1*). Aster-C global knock out mice were obtained from MMRRC (C57BL/6N-Atm1Brd Gramd1ctm1a(KOMP)Wtsi/JMmucd). The mutation was induced by insertion of the L1L2_Bact_P cassette at position 43812106 of Chromosome 16 upstream of the critical exon (exon 11). Aster-B $^{fl/fl}$ mice were generated by C57BL/6 ES cell-based gene targeting. Briefly, Exons 3-6 of the macrophage variant (exons 6-9 of intestine specific variant) were selected as conditional knockout region. To engineer the targeting vector, homology arms and conditional KO region were generated by PCR using BAC clone RP23-366E16 and RP23-184D8 from the C57BL/6 library as template. In the targeting vector, the Neo cassette was flanked by SDA (self-deletion anchor) sites. Aster-C $^{fl/fl}$ mice were generated from C57BL/6N-Atm1Brd Gramd1ctm1a(KOMP)Wtsi/JMmucd. The cassette is composed of an FRT site followed by lacZ sequence and a loxP site. This first loxP site is followed by a neomycin resistance gene under the control of the human beta-actin promoter, SV40 polyA, a second FRT site and a second loxP site. A third loxP site is inserted downstream of exon 11 at position 43812967. Exon 11 is thus flanked by loxP sites. The floxed allele was created by flp recombinase expression in mice carrying this allele. Aster-B $^{fl/fl}$, Aster-C $^{fl/fl}$, and Aster-B $^{fl/fl}$; Aster-C $^{fl/fl}$ were backcrossed into C57BL/6N background and crossed with Villin-CreERT2 mice to generate tamoxifen inducible intestine-specific knockout. After consecutive intraperitoneal injections of 1 mg tamoxifen in 100 µl corn oil for 5 days, intestine-specific knock out were generated. 3×HA Aster-B knock-in mice were generated on a C57BL/6N background by inserting a 3×HA tag into the first exon of Aster-B (*Gramd1b*) gene using a CRISPR/Cas9 based strategy. NPC1L1 knock out mice were a gift from Bao Liang Song, Wuhan University. They were also backcrossed to 3×HA Aster-B knock-in mice. For the *in vivo* kinetic absorption of radiolabeled cholesterol in the presence of ezetimibe or AI-3d, 8 weeks old female mice (C57BL/6J) were purchased from The Jackson Laboratory.

For gene expression, protein expression, and lipidomic analyses small intestines were excised and flushed with 50mL of ice-cold phosphate buffered saline (PBS) supplemented with protease inhibitor cocktail (Roche) and with 1mM dithiothreitol (DTT). Small intestines were cut into three segments with length ratios of 1:3:2 (corresponding to duodenum, jejunum, and ileum), and each segment was open longitudinally and scraped with ice-cold glass to isolate scrapings containing intestinal epithelial cells. Intestinal scrapings were snap frozen in liquid nitrogen and stored at -80°C or fixed in 10% formalin for histological analysis. Unless differently specified, duodenum was used for protein detection, proximal jejunum for lipidomic analysis and distal jejunum for gene expression analysis. Blood was collected by cardiac puncture (terminal) or by retro-orbital bleeding, and the plasma was separated by centrifugation. Unless, differently specified, mice were fed chow diet ad libitum and housed in a pathogen-free animal facility at 22°C, with a daylight cycle from 06:00 to 18:00. For western diet high cholesterol studies, male mice were fed a western diet containing 1.25% cholesterol (Research Diet, D09062501Ni) for 21 days. At the end of the study mice were fasted for 4hr before the euthanasia. Male mice were also used to investigate the effects of the NPC1L1 inhibitor ezetimibe on mice with deletion of Asters. In this regard, mice were fed for 3 days with a control diet containing 830mg/kg cholesterol or with a matched diet containing 0.01% ezetimibe. Mice were fasted overnight and refed for 2hr before the euthanasia. All animal experiments were approved by the UCLA Institutional Animal Care and Research Advisory Committee.

Histology and Immunohistochemistry

For histology, fragments of jejunum were fixed overnight in 10% neutral-buffered formalin at room temperature, embedded in paraffin and sectioned. Sections were deparaffinized and stained with H&E. For HA-Aster-B immunohistochemistry, 3×HA Aster-B were fed for 3 days with Ctrl diet or EZ diet and were treated three times (every 12 h) by oral gavage with 40mg/kg GW3965. After the last administration of GW3965, mice were fasted for 4 h and then orally administered with vehicle or 10 mg/kg ezetimibe 30 minutes before receiving a gastric gavage with corn oil only, or corn oil containing 6 mg of cholesterol. After 1 h mice were euthanized and perfused with 10ml cold HBSS. Small intestines were dissected and intestinal lumen was immediately flushed with 20ml cold HBSS to fully remove luminal content, followed by slow perfusion with 10ml of cold neutral buffered formalin for immediate fixation of the villi. Small intestine was longitudinally opened and Swiss-rolling of the intestine was performed from duodenum to distal ileum, with villus tip facing radially outwards. Intestine Swiss roll was then immobilized with foam biopsy pads in histology macro-cassette and fixed for 48 hours in formalin followed by dehydration in 70% ethanol. Tissue was paraffin embedded and 5μm cross-section was performed to reveal the entire roll face. Sections were deparaffinized and subjected to antigen retrieval with 10 mM sodium citrate (pH 6.0) in a sub-boiling water bath for 20 minutes. Slides were then incubated with the primary antibody overnight at 4°C. For IHC staining, slides were further incubated with Envision+ System-HRP Labelled Polymer Anti-Rabbit (Agilent, # K400311-2) at room temperature for 1 h and developed with ImmPACT DAB Peroxidase (HRP) Substrate (Vector, #SK4105).

Cholesterol absorption assays

For the acute cholesterol uptake assay, female mice were fasted for 4hr and then gavaged with 2μCi [14C] cholesterol (Perkin Elmer) in 200 μl olive oil. 2 hr later, mice were euthanized and small intestine was excised (between the base of the stomach and the cecal junction), flushed with 0.5 mM sodium taurocholate in PBS, and cut it into 2-cm segments, as previously described (*2*). Segments were incubated with 500 ml of 1N NaOH at 65°C overnight and mixed with ScintiSafe (Fisher Scientific), and scintillation was counted. Plasma and liver were also harvested, and liver bigger lobes were incubated with 1mL of 1N NaOH at 65°C overnight and mixed with ScintiSafe for scintillation counting.

For acute fatty acid uptake assay, the mice were fasted for 4hr and then gavaged with 2μCi [14C] triolein in 200 μl olive oil. The samples were processed as described above.

Cholesterol absorption was also measured in kinetic assays where mice were fasted for 10hr and then gavaged with 2μCi [14C] cholesterol in 200 μl olive oil. Blood was collected at time 0, 30 min, 1hr, 2hr, 4hr, 6hr and 12hr, and the plasma was separated by centrifugation. Radioactivity was measured by scintillation.

For studies in the presence of Poloxamer-407, 10gr of Poloxamer-407 were resuspended in 100mL of 0.9% NaCl saline, and stirred overnight at 4°C. 10mL/gr of body weight were administered by intraperitoneal injection right before the oil gavage with 2μCi [14C] cholesterol. Blood was collected at time 0, 1hr, 2hr, 3hr, and 4hr, and the plasma was separated by centrifugation. Radioactivity was measured by scintillation.

Fractional cholesterol absorption was measured by fecal dual isotope as previously described (*3*). Briefly, the mice were gavaged with 100 µl corn oil containing 0.5 µCi [14C] cholesterol µCi, 1 µCi [3H] sitostanol (American Radiolabeled chemicals) and 0.1 mg unlabeled cholesterol. Total fecal output was collected for 72 hr, snap frozen and pulverized using a pestle and a mortar. 500mg of pulverized feces were used for lipid extraction (*4*) with 19 volumes of 2:1 (v/v) chloroform:methanol at 60°C for 3min. The insoluble material was pelleted by centrifugation (1000g for 5min at 4°C). The chloroform:methanol extraction was repeated on the insoluble material. Solvent was removed by drying under N2 gas. Extracted lipids were saponified with 3mL of 1:1 (v/v) methanol:2N NaOH(aq) for 60min in a 60°C water bath. The neutral sterols were isolated with 3 sequential petroleum ether extractions, by adding 3mL of petroleum ether to each tube, mixing vigorously and separating the phases by centrifugation (1000g for 5min at 4°C). The upper, organic, phase was transferred to a scintillation vial. The contents of the scintillation vials were dried under N2 prior to addition of scintillation cocktail. The radioactivity was quantified by liquid scintillation counting. Cholesterol absorption was calculated using the following equation: % cholesterol absorption = ([14C]/[3H] dosing mixture – [ 14C]/[3H] feces)/([14C]/[3H] dosing mixture) × 100.

Lipid measurement in plasma

Plasma lipids were measured by colorimetric assay with the Wako L-Type TG M kit, the Wako Cholesterol E kit. For fast protein liquid chromatography (FPLC) lipoprotein plasma samples (pools from 4-5 mice/group) were injected into a Superose 6 10/300 (GE Healthcare Life Sciences) FPLC column and fractions were collected for measurement of cholesterol by colorimetric assay.

Chylomicron isolation

To isolate chylomicrons, 0.5mL of plasma were overlaid with 0.5mL of saline. The samples were then centrifuged at 35,000 rpm in an Optima MAX-XP Ultracentrifuge for 60 minutes at 18°C. After centrifugation, the top layer was collected as the chylomicrons fraction (~100 µL).

For d4-cholesterol experiment, F/F and I-B/C KO mice were gavaged with 500µg of d4-cholesterol in 100µL of corn oil and injected with 10mL/gr Poloxamer-407. 3.5 hours post-gavage, blood was harvested by cardiac puncture and centrifuged at 12,000xg for 3min. Plasma was collected. 700µL of cold PBS+8.6mM EDTA were transferred in ultracentrifuge tube and carefully underlaid with 300µL of plasma. Samples were centrifuged in Beckman TLA-120.2 at 100,000 rpm at 10°C for 2 hours. Chylomicrons were removed from the top of the tubes with a metal spatula, and dissolved overnight at 4°C. 10µL aliquots were used for lipid extraction and shotgun lipidomics.

Lipid measurement in liver

Livers were homogenized by dounce homogenizer in lysis buffer (10 mM Tris-HCl pH 7.4, 150 mM NaCl, 1 mM EDTA, protease inhibitors) and Folch method (*5*) was used to extract lipids from 0.5-1 mg liver protein. Protein content was measured by Bicinchoninic Acid Assay Protein Assay Kit (Pierce).

Lipid measurement in jejunum

Female mice were fasted for 4hr and gavaged with 2µCi [14C] cholesterol (Perkin Elmer) in 200 µl olive oil. 2hr later, mice were euthanized and small intestine was excised and flushed with 0.5

mM sodium taurocholate in PBS. Proximal jejunum (first 6 cm) was isolated, cut longitudinally, and minced with scissors. Then the minced tissue was and homogenized in 1.6 mL lysis buffer (10 mM Tris-HCl pH 7.4, 150 mM NaCl, 1 mM EDTA, protease inhibitors) using a dounce homogenizer. The jejunum homogenates were collected into a glass tube and lipids were extracted by Bligh and Dier method (*6*). In brief, homogenizer was washed with 2mL of methanol twice and the 4 mL were collected into the glass tube. 2mL of chloroform were added to the tube, which was vortexed for 30sec. Then 2mL of chloroform and 2mL 0.9% KCl were added to the tube, and the solution was vortexed for 15sec. Samples were centrifuged 2500rpm for 10min at room temperature. The lower layer was collected into a new glass tube, and the upper layer was subjected to an additional lipid extraction with 4mL of chloroform. Samples were centrifuged 2500rpm for 10min at room temperature, and the upper organic phase was collected into the same glass tube of the first extraction. The solvent was evaporated under N2 and lipid film was resuspended in 100mL of chloroform. Cholesterol and cholesterol esters were separated by thin-layer chromatography (TLC) on silica plates using the solvent system heptane: isopropyl ether: acetic acid (60: 40: 4). The silica plates were exposed on a phosphor screen (Cytiva/GE Healthcare Life Sciences) and the exposed screens were scanned with a bio-imaging analyzer (Typhoon Variable Mode Imager, GE, Piscataway, NJ) to quantify the incorporation of radioactivity in cholesterol and cholesterol esters.

Isolation of intestinal epithelial cells

Isolation of enterocytes in a gradient along the crypt/villus axis was performed as previously described (*7*). Briefly, after dissection, small intestine portions were cut longitudinally to efface intestinal mucosa and sliced it in pieces of 5-6 cm. Intestinal pieces were placed in tubes containing 20 ml of citrate buffer (27 mM sodium citrate, 96 mM sodium chloride, 1.5 mM potassium chloride, 8 mM potassium phosphate dibasic, 5.6 mM sodium phosphate dibasic, 1 mM DTT, pH 7.3) and incubated at 37°C for 5 min. Jejunal segments were then transferred into tubes contained 25 mL of prewarmed Enterocyte isolation Buffer ($Ca^{2+}$ and $Mg^{2+}$-free PBS with 1.5 mM EDTA and 1 mM DTT, pH 7.3) and shake at 37°C. Cell-containing supernatants from eight sequential fractions were collected after incubation for 11, 17, 23, 29, 35, 45, 60, and 80 min. After each time point, the jejunal pieces were transferred into new tubes with 15 ml of fresh isolation buffer. Residual cell suspension was strained (100uM), centrifuged at 1000g 5 min RT, and pellets were snap frozen in liquid nitrogen and put the pellet at -80. Fractions 1-3 were pooled to obtain the upper villus, fractions 4-5 were pooled into the middle villus, and fractions 6-8 were pooled to obtain the low villus/crypt fraction.

Protein isolation and immunoblot analysis

To prepare whole cellular lysates, enteroids from 4 wells were pooled and pelleted by quick spinning, and washed 5 times with 1mL of PBS. The pellets were resuspended in RIPA buffer supplemented with protease and phosphatase inhibitor cocktails (Roche), by harsh pipetting and incubation on rotation at 4°C for 30 minutes. To prepare membrane-enriched lysates from intestine, duodenal or jejunal scrapings were dounced in 500mL-1mL of lysis buffer (150mM NaCl, 1.5mM DTT, 50mM Tris HCl pH 7.4, 1.25mM EDTA, 0.1mM PMSF) containing protease and phosphatase inhibitor cocktails (Roche). Samples were quickly sonicated (3 pulses, 3 seconds on, 50%), and centrifuged at 4°C, 2000g for 10 minutes. Supernatant were collected and ultracentrifuged at 4°C, 100,000g for 45 minutes. Pellets were resuspended in the lysis buffer (100-150uL) using Hamilton syringe. Protein content was determined by the Bicinchoninic Acid Assay Protein Assay Kit (Pierce) and an equal amount of protein was loaded into the NuPAGE 4-12% Bis-Tris gels (Invitrogen). Proteins were transferred to PVDF blotting membrane (Amersham™ Hybond™, 10600023). Membranes were blocked for 1 hour at room temperature with 5% milk in TBS + 0.1% Tween 20 (TBST), and incubated with primary antibodies as indicated in Table S1. Horseradish peroxidase-conjugated anti-mouse, anti-goat and anti-rabbit IgG (Jackson) were used as secondary antibodies. The immune signal was visualized using the ECL kit (Amersham Biosciences).

RNA extraction and gene expression analyses

Total RNA was extracted using TRIzol (Invitrogen) and quantified by NanoDrop™ (Thermofisher) and reverse transcribed. cDNA was quantified by real-time PCR using iTaq Universal SYBR Green Supermix (Bio-Rad) on a QuantStudio 6 Flex 384-well qPCR system (Applied Biosystems). Gene expression levels were determined by using a standard curve. Each Housekeeping gene 36b4 was used for normalization and every sample was analyzed in duplicate. Primers used for real-time PCR are available upon request. For RNA sequencing, total RNA was extracted using TRIzol and the RNeasy Mini Kit with on-column DNase I digestion (Qiagen). Libraries were prepared using NEBNext Ultra II RNA Library Prep Kit for Illumina (New England Biolabs, Ipswich, MA, USA) and sequenced on Illumina HiSeq 4000 as paired-end 150 base pairs. Adapter and quality trimming of raw FASTQ files was performed using Trimmomatic. FastQC was used to analyze FASTQ files before and after trimming. Trimmed FASTQ files were aligned to GRCm38/mm10 using STAR. Alingned reads were visualized using Integrative Genomics Viewer (IGV). Gene counts were normalized and differential gene expression analysis was performed using DESeq2. Heatmap was created using ClustVis web tool.

Lipidomic analysis

Intestinal scrapings (approximately 100 mg of proximal jejunum scrapings) were collected in a 2 mL homogenizer tube pre-loaded with 2.8mm ceramic beads (Omni #19-628). PBS (0.75 mL) was added to the tube and the sample was homogenized in an Omni Bead Ruptor Elite (3 cycles of 10 seconds at 5 m/s with a 10 second dwell time). Homogenate containing 2-6 mg of original tissue was transferred to a glass tube for extraction. A modified Bligh and Dyer extraction (*8*) was carried out on all samples. Prior to biphasic extraction, an internal standard mixture consisting of 70 lipid standards across 17 subclasses was added to each sample (AB Sciex 5040156, Avanti 330827, Avanti 330830, Avanti 330828, Avanti 791642). Following two successive extractions, pooled organic layers were dried down in a Thermo SpeedVac SPD300DDA using ramp setting 4 at 35 degrees C for 45 minutes with a total run time of 90 minutes. Lipid samples were resuspended in 1:1 methanol/dichloromethane with 10mM ammonium acetate and transferred to robovials (Thermo 10800107) for analysis. Samples were analyzed on the Sciex 5500 with DMS device (Lipidyzer Platform) with an expanded targeted acquisition list consisting of 1450 lipid species across 17 subclasses (or the original acquisition list of 1100 lipids across 13 subclasses). Differential Mobility Device on Lipidyzer was tuned with EquiSPLASH LIPIDOMIX (Avanti 330731). Data analysis was performed on an in-house data analysis platform comparable to the Lipidyzer Workflow Manager (*9*). Quantitative values were normalized to mg of tissue, protein content, or plasma volume.

For cholesterol measurements, d7-cholesterol standard was added to each sample (Avanti 700041). Following extraction and lipidomics measurement, samples were derivatized with acetyl chloride (Fisher AA43262AD) and cholesterol was measured as acetyl esters as previous described (*10*).

In vivo back scattered electron (BSE) microscopy and NanoSIMS

2 WT and 2 Aster BC KO mice were fasted overnight, then refed for 2h in order to synchronize digestive status. In the meantime, a mixture containing 40 mg [13C] mixed fatty acids (Cambridge Isotopes Laboratories inc. Andover, MA) and 4mg [2H] cholesterol per mouse was generated by combining 40uL [13C] mixed fatty acids (1 mg/uL) with sunflower oil (10uL/mouse). Next 4mg [2H] cholesterol (dissolved in pure ethanol) was mixed into the solution by pipetting. Ethanol was removed by evaporation under a constant stream of nitrogen for approximately 1h. After 2h of refeeding, mice were orally gavaged with the fatty acid/cholesterol mixture. 2h later, mice were anesthetized then the abdominal cavity was opened and intestines were placed directly into fixative (2.5% glutaraldehyde, 2% paraformaldehyde, 2.1% sucrose, 0.1M sodium cacodylate) then, small, ring-shaped slices were cut from the appropriate regions of the small intestine. Intestine slices were then fixed in the same fixative at 4ºC overnight. The following day, the samples were washed 5x5min in cold 0.1M sodium cacodylate and then postfixed with reduced osmium (2% OsO4, 1.5% potassium ferricyanide, 0.1M sodium cacodylate) for 1h. Next, samples were washed 5x5min in cold H2O then treated with 1% thiocarbohydrazide for 20min at room temperature. Samples were then rinsed with H2O 5x5min and stained again with osmium (2%OsO4 in $H_2O$) for 30min. After another round of $H_2O$ washes, the tissue was stained with 1% uranyl acetate (SPI Chem.) at 4ºC overnight. Tissues were dehydrated by incubating in a graded series of ethanol solutions (30%, 50%, 70%, 85%, 95%, 100%X3) 8min each. Then tissues were infiltrated with 33% (in acetone) EMbed812 for 1h, 66% overnight, then 100% for 4h. Individual tissue pieces were then embedded in a flat mold and polymerized in a vacuum oven for 48h. Resin blocks were then trimmed and 500 nm sections were collected onto small silicon wafers using a Leica UC6 ultramicrotome and a Diatome diamond knife. For backscattered electron microscopy: Wafers were then mounted onto SEM stubs with double-sided carbon tape and imaged using a Zeiss Supra 40VP SEM equipped with a backscattered electron detector. Accelerating voltage was set to 13KeV and working distance at approx. 5mm. BSEM was performed at the Electron Imaging Center for Nanomachines which is a part of the California NanoSystems Institute at UCLA. For correlative backscattered electron microscopy and NanoSIMS imaging, regions of interest were imaged using an FEI Verios XHR SEM with the 2-kV electron beam of 0.2-nA beam current and a backscattered electron detector (CBS). Mosaics of images were collected with the Maps 2.0 software and were stitched with Grid/Collection stiching in Image J (Preibisch et al., bioinformatics 2009). The same samples were coated with 5 nm platinum on the surface and transferred into a NanoSIMS 50L (CAMECA, France) for NanoSIMS analysis. A 133Cs+ primary ion beam of ~1 nA beam current was used for the implantation of caesium onto the sample surface (aperture D1 = 1) to reach a total ion dose of ~1×1017 ions/cm$^2$. 1H–, 2H–, 12C–, and 13C– secondary ion signals were collected to visualize and quantify the distributions of [2H]cholesterol and 13C-labelled mixed fatty acids. Images were obtained using aperture D1 = 2 with a current of ~3 pA, a dwell time of 3.0 ms/pixel, and a raster size of 30 µm × 30 µm. Mosaics of NanoSIMS images were obtained on the same areas that were mapped with backscattered electron imaging.

Isolation of murine crypts

Murine crypts were isolated from the jejunums of 8-12 week old 3XHA Aster-B, WT, and Aster BC KO mice as previously described (*11*). Animals were euthanized with isoflurane and the jejunum was harvested and flushed with ice-cold PBS. The intestine was inverted to expose the villous surface and submerged in 30 ml 2.5mM EDTA/PBS on a rocker at 4°C for 30 minutes. The supernatant was discarded and replaced with 15 mL PBS. The tube was then vortexed in 3 second pulses, 10 times, to release crypts from the tissue. Leaving the tissue behind, the mixture was poured into a new 15 mL conical tube and settled on ice for 5 minutes to allow unwanted cells and connective tissue to fall to the bottom. The supernatant containing the crypt fraction was filtered through a 70 mm cell strainer into a new 15 ml conical tube. The crypts were centrifuged at 100g for 2 minutes. The pellet was washed with 5 mL PBS and centrifuged at 100g for 2 minutes. The supernatant was removed and the pellet was suspended in Matrigel to a final concentration of 200 crypts per 25 µl. 25 µl of crypt/Matrigel suspension was plated into the wells of a 48 well plate and placed in a 37°C, 5% CO2 incubator for 20 minutes to polymerize. 250µl of growth media was added to each well. Growth media contained the following: 50% Advanced DMEM/F12 (Gibco 12634-010), 50% L-WRN conditioned media (*12*) , 2mM GlutaMAX, 10 mM HEPES, 100 U/ml penicillin/100 µg/ml streptomycin, 1X N2 supplement (Gibco 17502-048), 1X B27 supplement (Gibco 17504-044), 50 ng/ml EGF (Peprotech 315-09), 1 mM N-acetylcysteine (Sigma A9165-5G), 2.5 µg/ml fungizone, and 200 µg/ml normocin. 10 µM Y-27632 (Sigma Y0503) was added for the first few days after extraction and after splitting.

Culture and expansion of murine enteroids

Enteroids were passaged every 7-10 days. After removing the media, the matrigel was broken with ice-cold PBS to release the enteroids. The solution was pelleted and washed 2 additional times with PBS. After the final wash, the pellet was suspended in 200 µl of PBS and the enteroids were broken by pipetting 5-10 times. The broken enteroids were centrifuged and the supernatant was aspirated. 10µl of growth media was added to the pellet before resuspending in matrigel. 25 µl of the enteroid/matrigel suspension was plated into the wells of a 48 well plate and growth media was added to the wells after polymerization. Enteroids passaged more than 4 times, but less than 30 times, were utilized for the experiments. Prior to use, enteroids were differentiated for 5 days in murine differentiation media, containing Advanced DMEM/F12, 2 mM GlutaMAX, 10 mM HEPES, 100U/ml penicillin/100µg/ml streptomycin, 1X N2 supplement, 1X B27 supplement, 50 ng/ml EGF, 50 ng/ml noggin (Peprotech 250-38), 50 ng/ml r-Spondin1 (R and D 3474-RS), 10 µM Y-27632, 2.5 µg/ml fungizone, and 200 µg/ml normocin.

Culture of apical-out murine enteroids

Basolateral out enteroids were everted to generate apical-out enteroids, as previously described (*13*). The matrigel of young basolateral-out enteroids (day 3-5) was broken with ice cold PBS and transferred into a microcentrifuge tube. The enteroids were spun down and the supernatant was aspirated. To ensure adequate removal of matrigel, the pellet was resuspended in cell recovery solution (Corning 354253) and incubated at 4°C for 20 minutes. The enteroids were pelleted and resuspended in 1 ml of 5 mM EDTA/PBS solution. The solution was transferred to a 15 ml conical tube containing 10-12 ml 5 mM EDTA/PBS and incubated on a rotating platform at 4°C for 1 hour. The tubes were centrifuged at 300g for 3 min and the ETDA/PBS solution was aspirated. The pellet containing the enteroids was washed with 5 ml PBS and centrifuged at 300g for 3 min. The pellet was resuspended in enteroid growth media with 10 µM Y-27632 and plated in ultra-low attachment plates. After 1-2 days, the media was exchanged with murine differentiation media with 10 µM Y-27632 and the enteroids were allowed to differentiate for 5 days. The media was changed every 2-3 days.

Culture of basolateral out human enteroids

De-identified human enteroids were provided as a gift from the laboratory of Dr. Martin Martin. The enteroids were expanded and passaged every 7-10 days, using the same method for murine enteroids. Human enteroids were cultured in growth media, containing 50% Advanced DMEM/F12 (Gibco 12634-010), 50% L-WRN conditioned media (*12*), 2 mM GlutaMAX, 10 mM HEPES, 100 U/ml penicillin/100 μg/ml streptomycin, 1X N2 supplement, 1X B27 supplement, 50 ng/ml EGF, 1 mM N-acetylcysteine, 500 nM A83-01 (Tocris 909910-43-6), 10 μM SB202190 (Sigma S7067), 2.5 μg/ml fungizone, and 200 μg/ml normocin. 10 μM Y-27632 was added for the first few days after passaging the enteroids. Enteroids were differentiated for 5 days prior to use for experiments. Human differentiation media contained Advanced DMEM/F12, 2 mM GlutaMAX, 10 mM HEPES, 100 U/ml penicillin/100 μg/ml streptomycin, 1X N2 supplement, 1X B27 supplement, 50 ng/ml EGF, 50 ng/ml noggin, 50 ng/ml r-Spondin, 500 nM A83-01, 10 μM Y-27632, 2.5 μg/ml fungizone, and 200 μg/ml normocin.

Culture of human intestinal epithelial cells on transwell membranes for gene expression

Human intestinal epithelial cells (IEC) were derived from basolateral-out human jejunal enteroids to culture IEC's as a monolayer on transwell membranes. Basolateral-out enteroids were extracted from matrigel with PBS and incubated in cell recovery solution for 20 min at 4°C. The enteroids were pelleted and the supernatant was removed. The pellet was resuspended in TrypLE for 10 min at 37°C to obtain single cells. Media was added to stop the reaction and the mixture was centrifuged to pellet the cells. The cells were resuspended in growth media containing 50% Advanced DMEM/F12 (Gibco 12634-010), 50% L-WRN conditioned media, 2 mM GlutaMAX, 10 mM HEPES, 100 U/ml penicillin/100 μg/ml streptomycin, 1X N2 supplement, 1X B27 supplement, 50 ng/ml EGF, 1 mM N-acetylcysteine, 500 nM A83-01, 10 μM SB202190, 10 μM Y-27632, 2.5 μg/ml fungizone, and 200 μg/ml normocin. 150,000 cells were plated per transwell. On day 2, the media was changed to human differentiation media and changed every other day. After 5 days of differentiation, cells were deprived of cholesterol by overnight incubation with 1mM Ro 48-8071 (Cayman Chemical, 10006415) and 50mM mevalonate. Of note, human differentiation media contains no cholesterol. After 16 h, cells were pre-treated with vehicle or 5 mM AI-3d for 2.5 h. The monolayer was then loaded with mixed micelles containing 6mM oleic acid (Cayman Chemical, 90260), 0.5mM cholesterol (Sigma, C8667), 2mM 2-palmitoylglycerol (Cayman Chemical, 17882), 2mM Lyso-PC (Avanti, 845875C), 40mM sodium taurocholate (Sigma, 86339), 1mM Ro 48-8071, and 50mM mevalonate, in the presence of vehicle or 5mM AI-3d. After 4 h, cells were processed for RNA extraction.

Immunofluorescence

HA tagged Aster-B enteroids grown basolateral out were differentiated for 5 days and pre-treated with 1mM Ro 48-8071 and 50mM mevalonate for 1.5 h, then incubated with vehicle or 150 μM MbCD-cholesterol for 2h. The enteroids were extracted from matrigel using ice cold PBS and incubated in cell recovery solution for 1 hour at 4°C. The enteroids were washed with PBS, then fixed using 4% paraformaldehyde for 20 minutes at room temperature. The enteroids were allowed to settle to the bottom and the fixative was aspirated. Permabilization was performed with 0.3% Triton for 15 minutes at room temperature. The enteroids were blocked with 10% goat serum in PBS for 1 hour at room temperature and incubated in primary antibody (Anti-HA C29F4, 1:1000, CST 3724S) overnight at 4°C. The following day, the enteroids were washed with 2X PBS for 2 minutes, 3 times and incubated with secondary antibody (Anti-Rabbit AF594, 1:1000, CST 8889S) for 60-90 minutes at room temperature. The enteroids were washed again with 2X PBS for 2 minutes, 3 times; and then transferred to a glass slide. ProLong Diamond Antifade Mountant with DAPI (Invitrogen P36962) was applied to the slide prior to placement of the cover slip. The same protocol was applied to apical out enteroids with the exception of incubating in cell recovery solution. Imaging was performed on a Zeiss LSM 900 confocal microscope equipped with 405nm, 488nm, 561nm and 640nm laser lines using a Plan-Apochromat 20x/0.8 or Plan-Apochromat 40x/1.2 objective and Airyscan 2 GaAsP-PMT detector. Identical laser intensity settings were applied to each experimental sample set and Z-stacks were performed to acquire equivalent section thicknesses. After acquisition, a maximum intensity projection of the Z-stack was applied using ZEN Blue 3.0 software. HA tagged Aster-B enteroids grown apical out were processed in a similar fashion with the exception of a longer cholesterol depletion period with 1mM Ro 48-8071 and 50mM mevalonate for 16 h overnight. On the day of the experiment, enteroids were pre-treated with vehicle or 0.025 mM ezetimibe for 2 hours before treatment with mixed micelles containing 6mM oleic acid, 0.5mM cholesterol, 2mM 2-palmitoylglycerol, 2mM Lyso-PC, 40mM sodium taurocholate, 1mM Ro 48-8071, and 50mM mevalonate, in the presence of vehicle or 0.025 mM ezetimibe for an additional 2h. Enteroids underwent fixation and staining as outlined above.

Purification and fluorophore conjugation of ALOD4

pALOD4 was a gift from Arun Radhakrishnan (Addgene plasmids # 111026) (*14*) and was purified as previously described (*15*). Briefly, plasmid encoding ALOD4 was expressed in BL21 (DE3) pLysS E. coli (Invitrogen). Cell pellets were lysed by sonication in lysis buffer with 50mM NaH2PO4, pH 7.0, 300mM NaCl, 1mg/ml lysozyme, 1mM DTT, 2mM phenylmethylsulfonyl fluoride (PMSF), protease inhibitor cocktail tablet (Themo Scientific). Protein was bound to HisPur Ni-NTA Agarose resin (Thermo Scientific), washed twice with 50 mM imidazole, eluted with 300 mM imidazole, and subjected to size exclusion chromatography on Superdex 200. Fractions containing ALOD4 were concentrated to 1 mg/ml and stored at -80C with glycerol. ALOD4 was conjugated to Alexa 488 C5 maleimide (Thermo Fisher) followed by affinity chromatography using HisPur Ni-NTA Agarose resin as above. Samples were dialyzed to remove imidazole prior to determination of labeling efficiency on Nanodrop (Thermo Fisher).

ALOD4 staining in enteroids

Human and murine basolateral out enteroids were differentiated for 5 days. Enteroids were deprived of cholesterol in differentiation media containing 1mM Ro 48-8071 and 50mM mevalonate for 16 h. Murine WT and BC KO enteroids were treated with vehicle or 150 μM MbCD-cholesterol for 1h. For the AI3d experiment, murine and human WT enteroids were pre-treated with vehicle or 5mM AI-3d + 1mM Ro 48-8071 and 50mM mevalonate for 5h, and then treated with vehicle or 150 μM MbCD-cholesterol for 1h in presence or absence of 5 mM AI3d.

Enteroids in matrigel were washed with cold dPBS (+ Ca, + Mg) and 0.2% bovine serum albumin (BSA), 3 times. The enteroids were then collected in 400 μl of cell recovery solution with ALOD4 at a concentration of 20μg/ml and remained in solution for 1 hour at 4°C. After incubation, the enteroids were washed with dPBS (+ Ca, + Mg) and 0.2% BSA 4 times, 2 minutes each prior to fixation with 4% paraformaldehyde at room temperature for 15 minutes. The enteroids were again washed with dPBS (+ Ca, + Mg) and 0.2% BSA 4 times, 5 minutes each. The sample was mounted on glass slides with ProLong Diamond Antifade Mountant with DAPI and secured with a cover slip. Imaging was performed on a Zeiss LSM 900 confocal microscope using 405nm and 488nm laser lines for excitation. Images were acquired using a

Plan-Apochromat 20x/0.8 or Plan-Apochromat 40x/1.2 objective and Airyscan 2 GaAsP-PMT detector. Identical laser intensity settings were applied to each experimental sample set and Z-stacks were performed to acquire equivalent section thicknesses for comparison across treatments. After acquisition, a maximum intensity projection of the Z-stack was applied using ZEN Blue 3.0 software.

Caco-2

Caco-2 cells were seeded in 12wells transwell plate (Corning) at the density of $3 \times 10^5$ cells/cm$^2$ (approximately 350,000 cells/well) and cultured in DMEM + 10% fetal bovine serum (FBS) with 1% PenStrep (Gibco), on both apical and basolateral side. Cells were cultured in transwell up to 20 days, changing media every other day, and TEER was measured at day 7, day 14 and day 20. Cells with a TEER measurement around 1200 were considered ready for experiments. For AI-3d treatment, cells at day 16 of differentiation were cholesterol deprived by overnight incubation with DMEM containing 1% lipoprotein deficient serum (LPDS), 1mM Ro 48-8071 (Cayman Chemical, 10006415), and 50mM mevalonate. After 16 h, cells were pretreated with vehicle or 5mM AI-3d in DMEM with 1% LPDS, 1mM Ro 48-8071, and 50mM mevalonate for 2.5 h. Caco-2 cells were then loaded with DMEM supplemented with mixed micelles containing 6mM oleic acid (Cayman Chemical, 90260), 0.5mM cholesterol (Sigma, C8667), 2mM 2-palmitoylglycerol (Cayman Chemical, 17882), 2mM Lyso-PC (Avanti, 845875C), 40mM sodium taurocholate (Sigma, 86339), 1mM Ro 48-8071, and 50mM mevalonate, in the presence of vehicle or 5mM AI-3d. After 3 h, cells were processed for RNA extraction.

McArdle

CRL-1601 (a McArdle RH7777 rat hepatoma cell line) cells were maintained in DMEM containing 100 units/ml penicillin and 100 µg/ml streptomycin sulfate supplemented with 10% FBS. For cholesterol depletion cells were incubated with depleting medium (DMEM containing 5% LPDS, 10 µM compactin, 50 µM mevalonate, and 1.5% MbCD) for 1 h. For cholesterol-replenishment, cells were incubated in DMEM containing 5% LPDS, 10 µM compactin, 50 µM mevalonate, and 100µM MbCD-cholesterol, for 2h. Stable cell lines expressing NPC1L1-EGFP and HA-Aster-B were selected with 1 µg/mL puromycin.

Protein Expression and Purification

Protein expression and purification of mouse Aster-$C_{296\text{-}517}$ was undertaken as described previously for Aster-A (*1*). Excess EZ was added to partially purified Aster-C (molar ratio 1:5, protein: ligand) from a 10 mg/ml stock solution dissolved in ethanol before loading the complex on a Superdex S-200 column. The fractions containing the Aster-C: EZ complex were concentrated to 6 mg/ml and used for the crystallisation experiments.
Site directed mutagenesis was performed to generate the following mouse Aster-$A_{334\text{-}562}$ mutants: AsterA_L400A_F405I, AsterA_L400A and AsterA_F405I, and the following mouse Aster-$C_{296\text{-}517}$ mutants: AsterC_A357L_I362F, AsterC_A357L and AsterC_I362F. The resulting inserts were cloned and purified as described previously (*16*), without adding any compound to the protein samples. Instead,10 % of glycerol was added to the gel filtration buffer and the resulting samples were used for circular dichroism experiments.

Crystallization and X-Ray Structure Determination

Crystals of the Aster-$C_{296\text{-}517}$ in complex with EZ ligand were obtained using sitting drop vapor diffusion at room temperature. Crystals were grown using 0.2 M sodium chloride, 0.1 M MES pH 6.5, and 10% PEG 4000 (condition B4, PROPLEX, Molecular Dimensions). The crystals were cryo-protected with 30% glycerol, 0.2 M sodium chloride, 0.1 M MES pH 6.5, and 10% PEG 4000. Data were collected to 1.6 Å on the I24 beamline at Diamond Light Source, UK. Data were processed using XDS (within Xia2) and Pointless/Aimless (within CCP4) (*17-19*). The structure was solved using molecular replacement using Phaser (within CCP4) (*20*) and the Aster-A structure (PDB ID code 6GQF) as a template (*1*). Model fitting and refinement were performed using Coot, ArpWarp, Refmac (within CCP4), and Phenix (*21-24*). The structure has been submitted to the PDB database, PDB ID code 8AXW.

<u>Circular Dichroism</u>

Thermal unfolding of mouse Aster-$A_{334-562}$ wild-type and mutants, Aster-$B_{303-533}$ and Aster-$C_{296-517}$ wild-type and mutants was monitored by Circular Dichroism (CD) spectroscopy over a range of 200 to 250 nm, using a Chirascan Spectrometer (Applied Photophysics) equipped with a temperature controller (Quantum Northwest TC125). CD spectra were measured from samples in 1-mm-path-length quartz cuvettes, using a scanning speed of 100 nm/min, a spectral bandwidth of 1 nm, and a response time of 1 s. The thermal denaturation or unfolding profile of the proteins was characterized by measuring the ellipticity changes at 222 nm induced by a temperature increase from 20 to 90 ºC with steps of 1 degree. Samples of 1 mg/ml protein, apo or in complex with 25-HC, EZ or U18666A, were used. Melting temperature values were obtained by analysing the data using GraphPad Prism and a nonlinear regression analysis.

**Table S1. Antibodies used for detection in immunoblot analyses**

| Target | Vendor | Cat. number | Dilution | Species |
|---|---|---|---|---|
| Aster-A | Pierce | Customized antibody | 1:1000 | Rabbit |
| Aster-B | Pierce | Customized antibody | 1:1000 | Rabbit |
| HA-tag (C29F4) | Cell Signaling | 3724S | 1:1000 for WB<br>1:1000 for IF<br>1:800 for IHC | Rabbit |
| His-Tag (27E8) | Cell Signaling | 2366S | 1:1000 for WB<br>1:400 For IF | Mouse |
| ApoB | Chemicon | AB742 | 1:7500 | Goat |
| HMGCR | Millipore | ABS229 | 1:1000 | Rabbit |
| FDPS | Abcam | ab153805 | 1:1000 | Rabbit |
| LDL-R | Cayman Chemical | 10007665 | 1:1000 | Rabbit |
| SQLE | Sigma | AV42101 | 1:1000 | Rabbit |
| CALNEXIN | Abcam | ab10286 | 1:2000 | Rabbit |
| TUBULIN | Abcam | ab15568 | 1:10000 | Rabbit |

| | |
|---|---|
| Wavelength | 0.9686 |
| Resolution range | 40.93 - 1.6 (1.657 - 1.6) |
| Space group | $P\ 4_1\ 2_1\ 2$ |
| Unit cell | 49.62 49.62 144.77 90 90 90 |
| Total reflections | 552,892 (27,706) |
| Unique reflections | 24,703 (2,330) |
| Multiplicity | 22.4 (11.9) |
| Completeness (%) | 99.6 (96.5) |
| Mean I/sigma(I) | 30.3 (2.5) |
| Wilson B-factor | 18 |
| R-merge | 0.295 (0.529) |
| R-meas | 0.302 (0.553) |
| R-pim | 0.061 (0.154) |
| CC1/2 | 0.976 (0.894) |
| CC* | 0.994 (0.972) |
| Reflections used in refinement | 24,702 (2,330) |
| Reflections used for R-free | 1,245 (129) |
| R-work | 0.1724 (0.1933) |
| R-free | 0.2059 (0.2365) |
| CC(work) | 0.961 (0.919) |
| CC(free) | 0.937 (0.919) |
| Number of non-hydrogen atoms | 1666 |
| macromolecules | 1433 |
| ligands | 140 |
| solvent | 168 |
| Protein residues | 171 |
| RMS(bonds) | 0.012 |
| RMS(angles) | 1.15 |
| Ramachandran favored (%) | 98.82 |
| Ramachandran allowed (%) | 1.18 |
| Ramachandran outliers (%) | 0.00 |
| Rotamer outliers (%) | 0.00 |
| Clashscore | 4.04 |
| Average B-factor | 23 |
| macromolecules | 22 |
| ligands | 33 |
| solvent | 32 |

**Table S2. Data collection and refinement statistics for the crystal structure of Aster-C with Ezetimibe.**

Statistics for the highest-resolution shell are shown in parentheses.

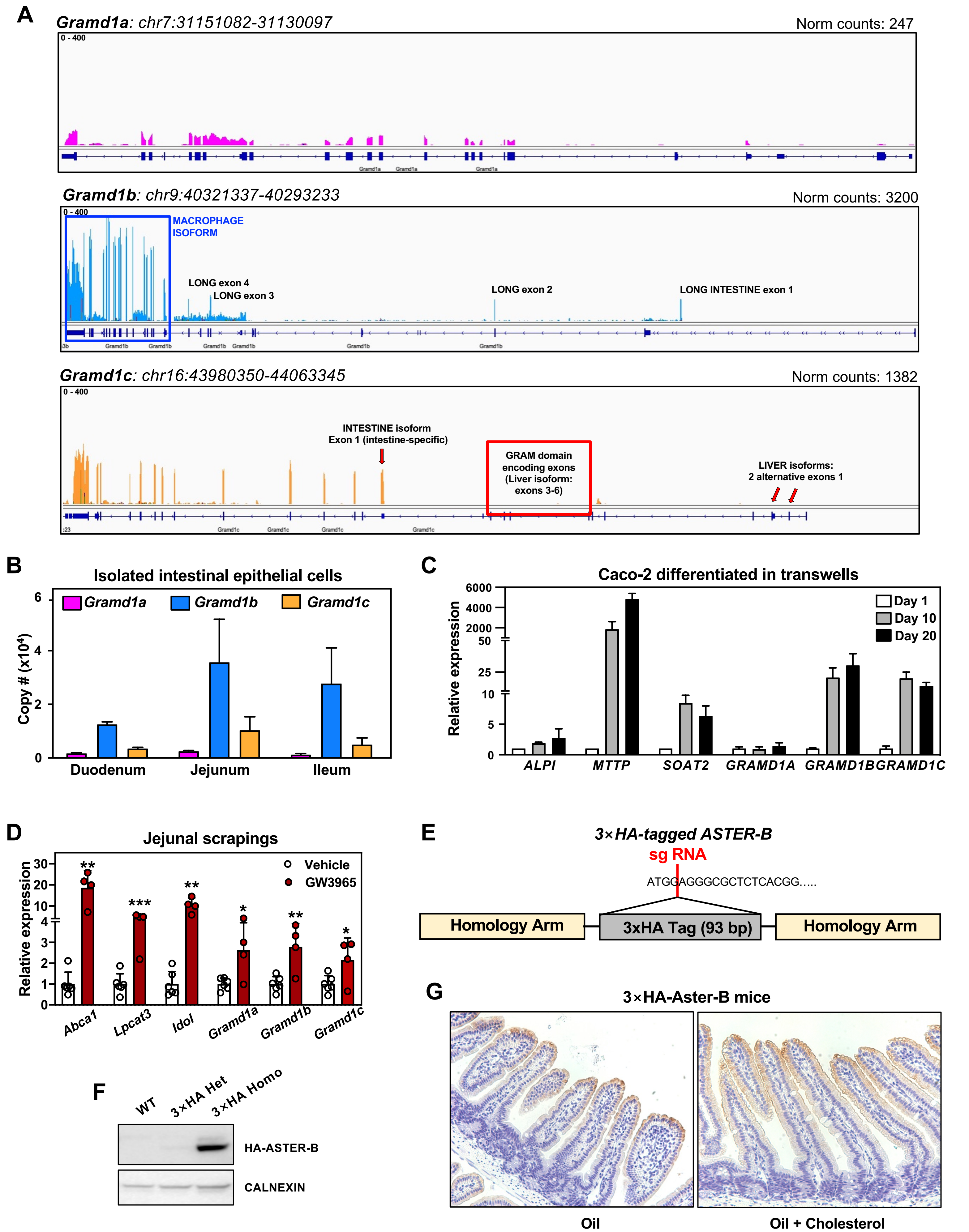


Ferrari et al, Fig. S1

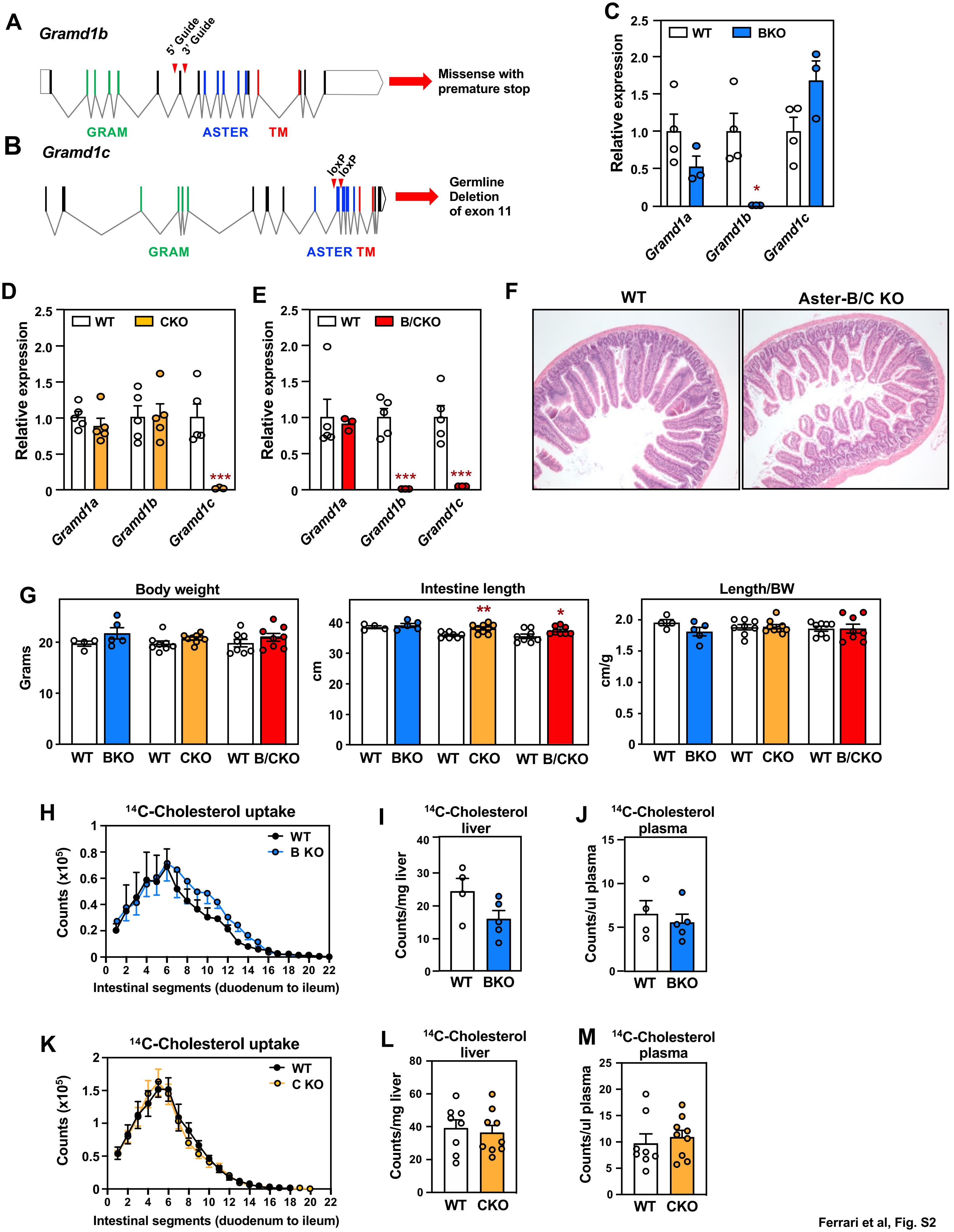


Ferrari et al, Fig. S2

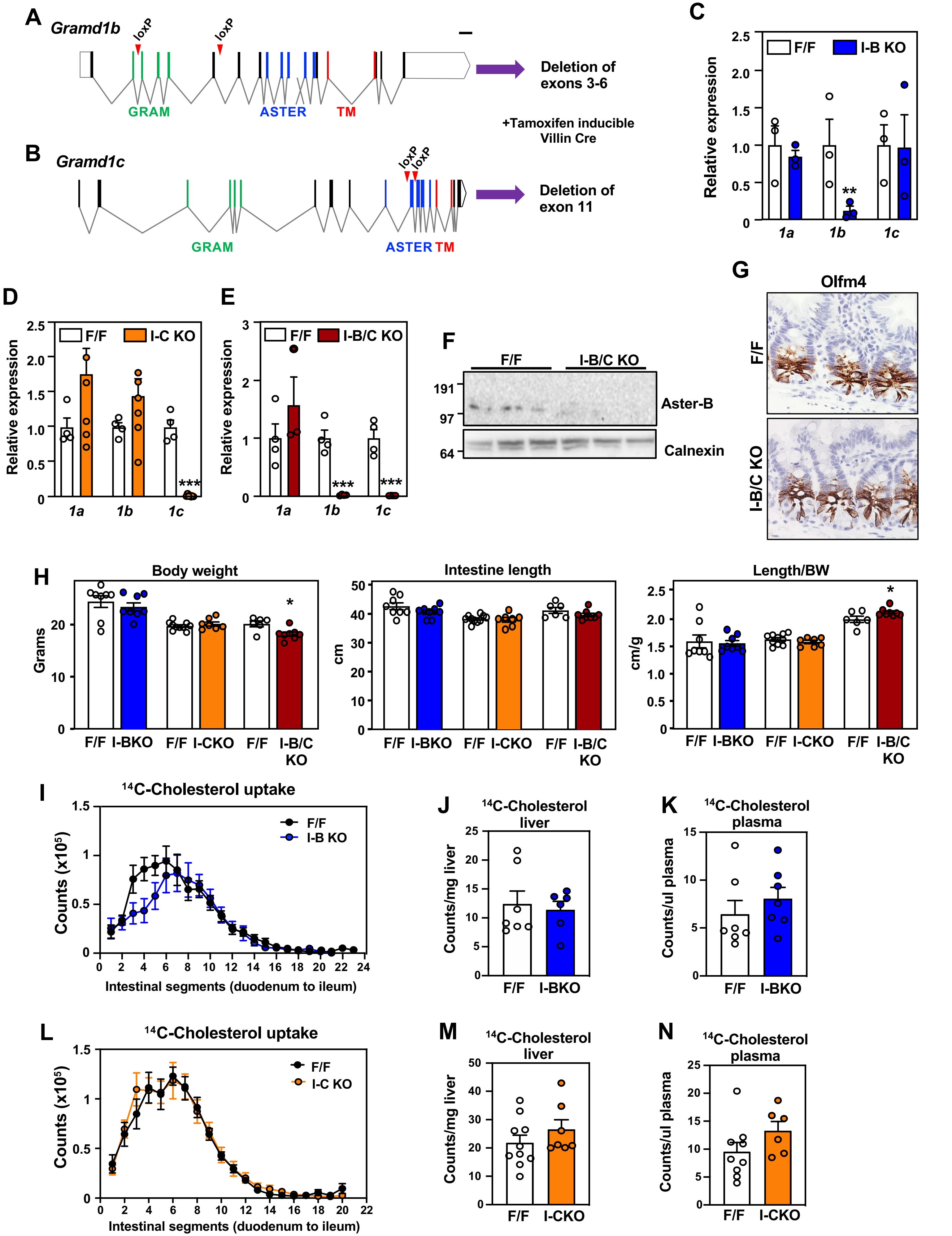


Ferrari et al, Fig. S3

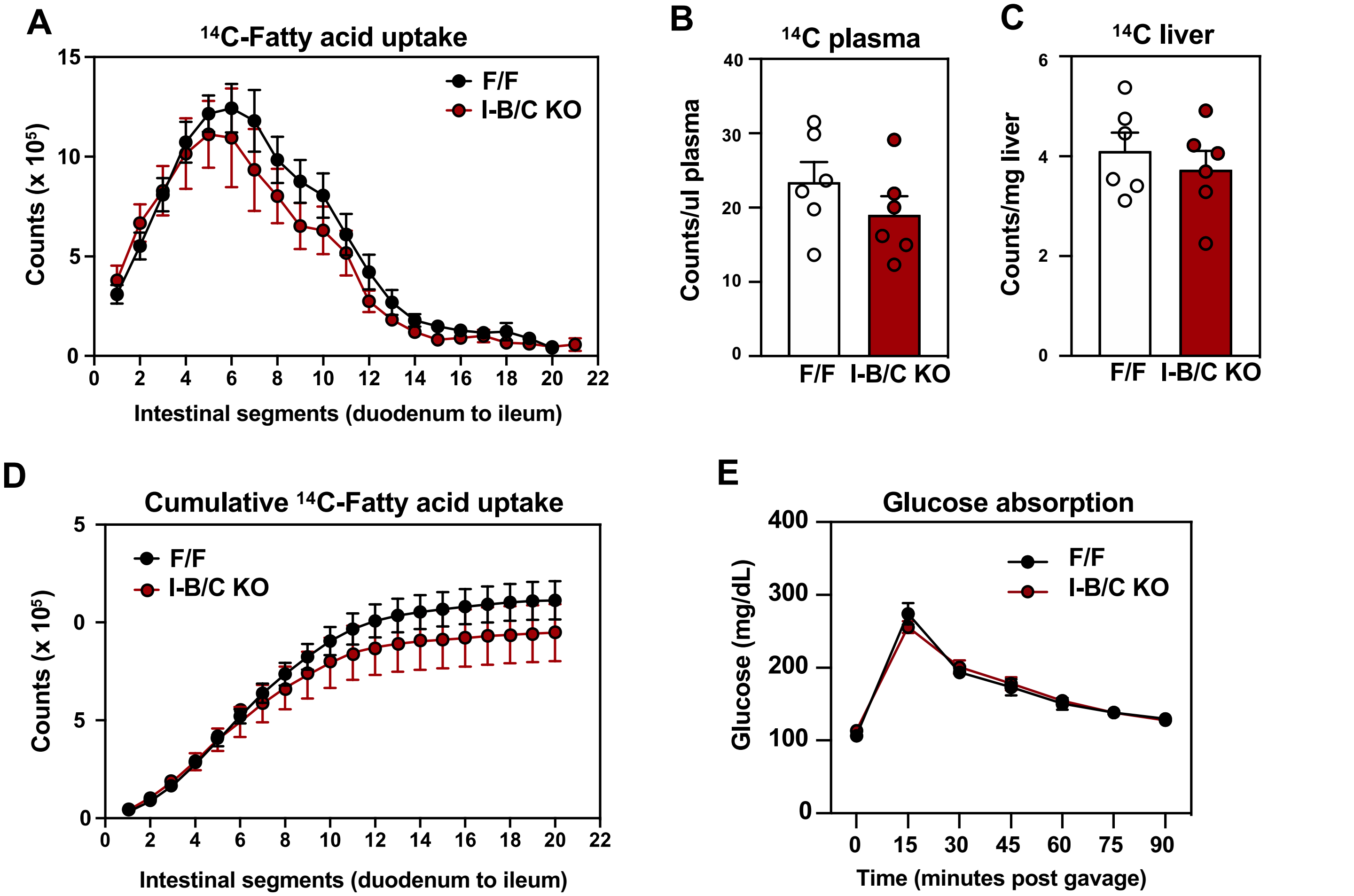
A
14C-Fatty acid uptake
Counts (x 10^5)
Intestinal segments (duodenum to ileum)
F/F
I-B/C KO
B
14C plasma
Counts/ul plasma
F/F
I-B/C KO
C
14C liver
Counts/mg liver
F/F
I-B/C KO
D
Cumulative 14C-Fatty acid uptake
Counts (x 10^5)
Intestinal segments (duodenum to ileum)
F/F
I-B/C KO
E
Glucose absorption
Glucose (mg/dL)
Time (minutes post gavage)
F/F
I-B/C KO

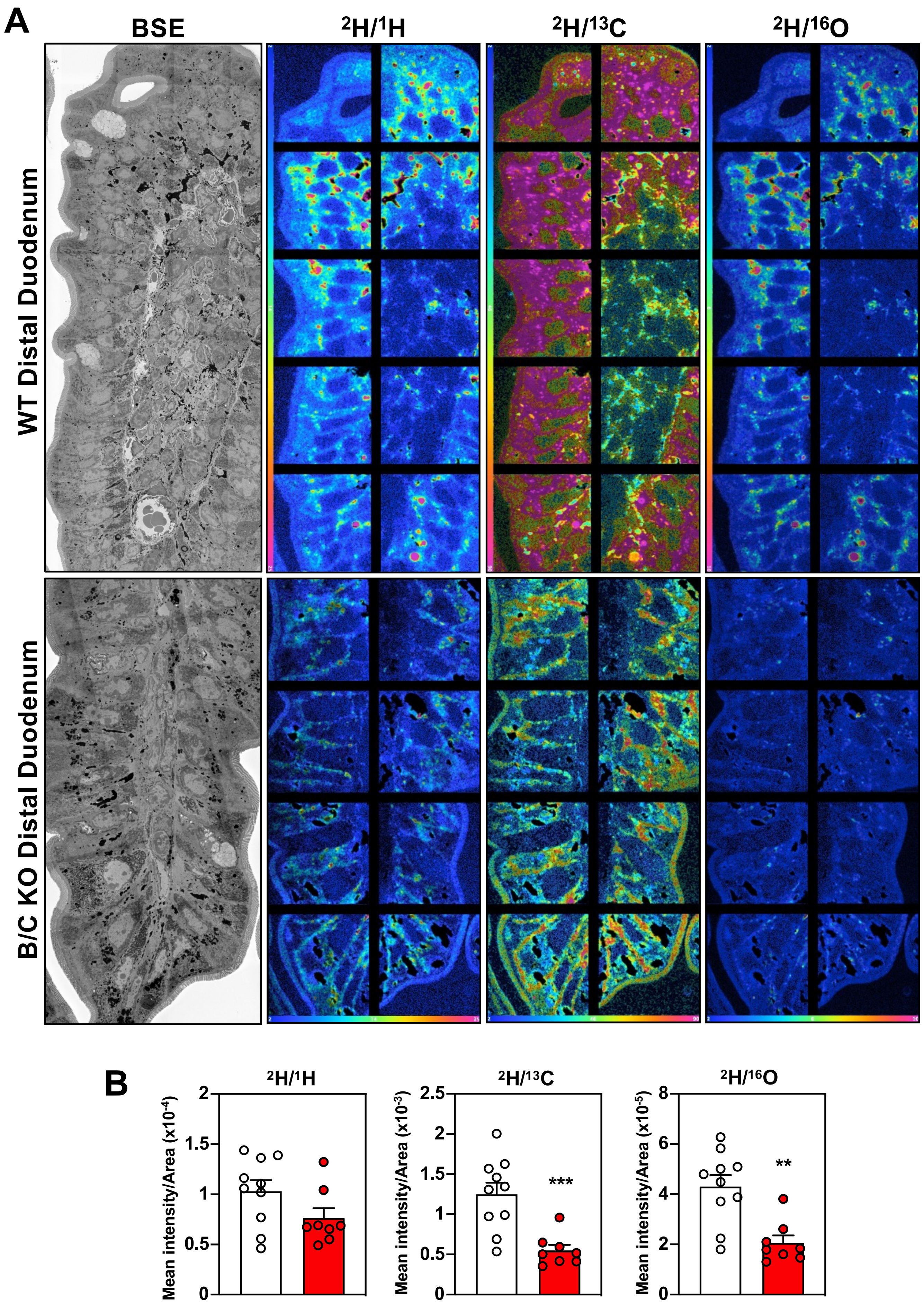


Ferrari et al, Fig. S5

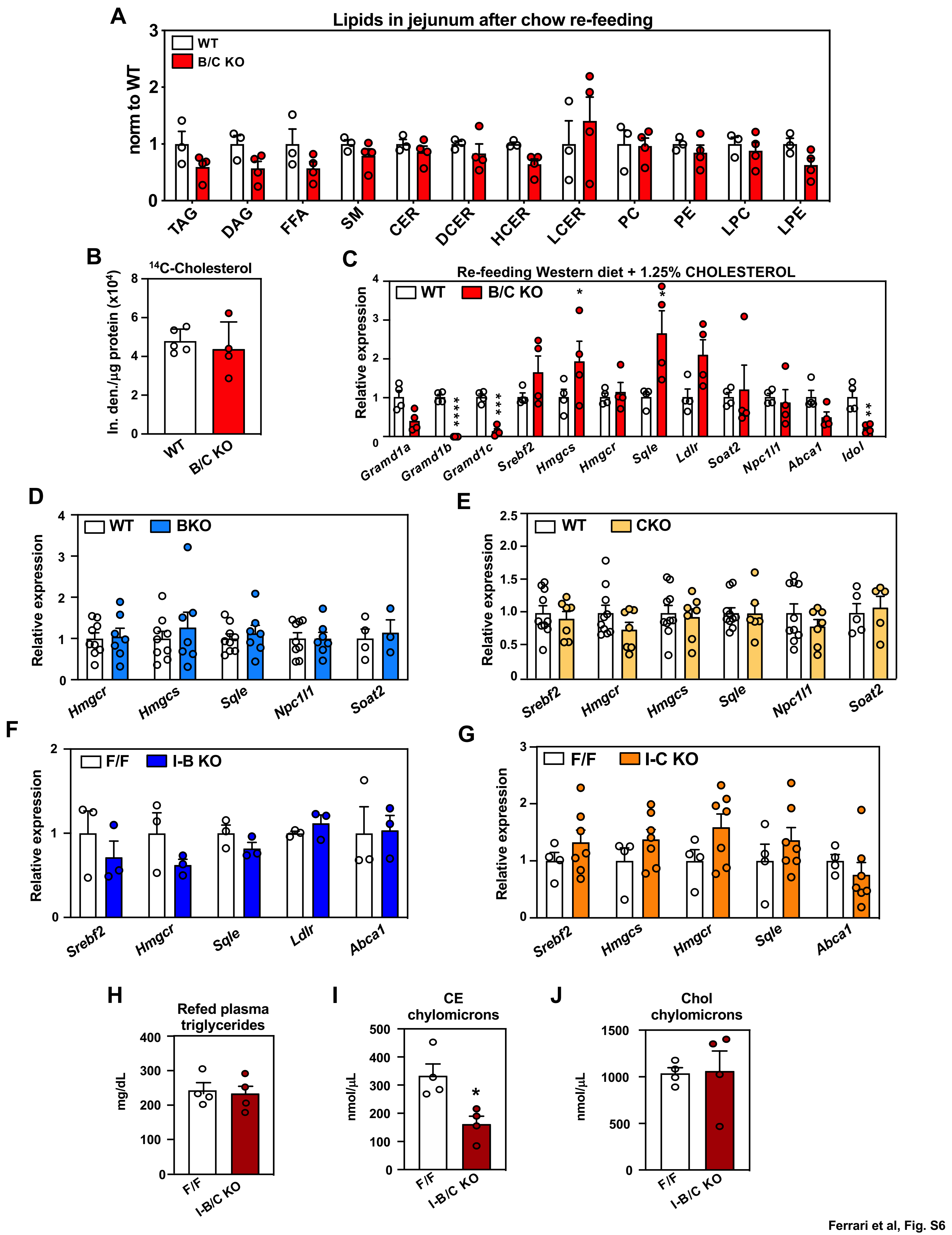


Ferrari et al, Fig. S6

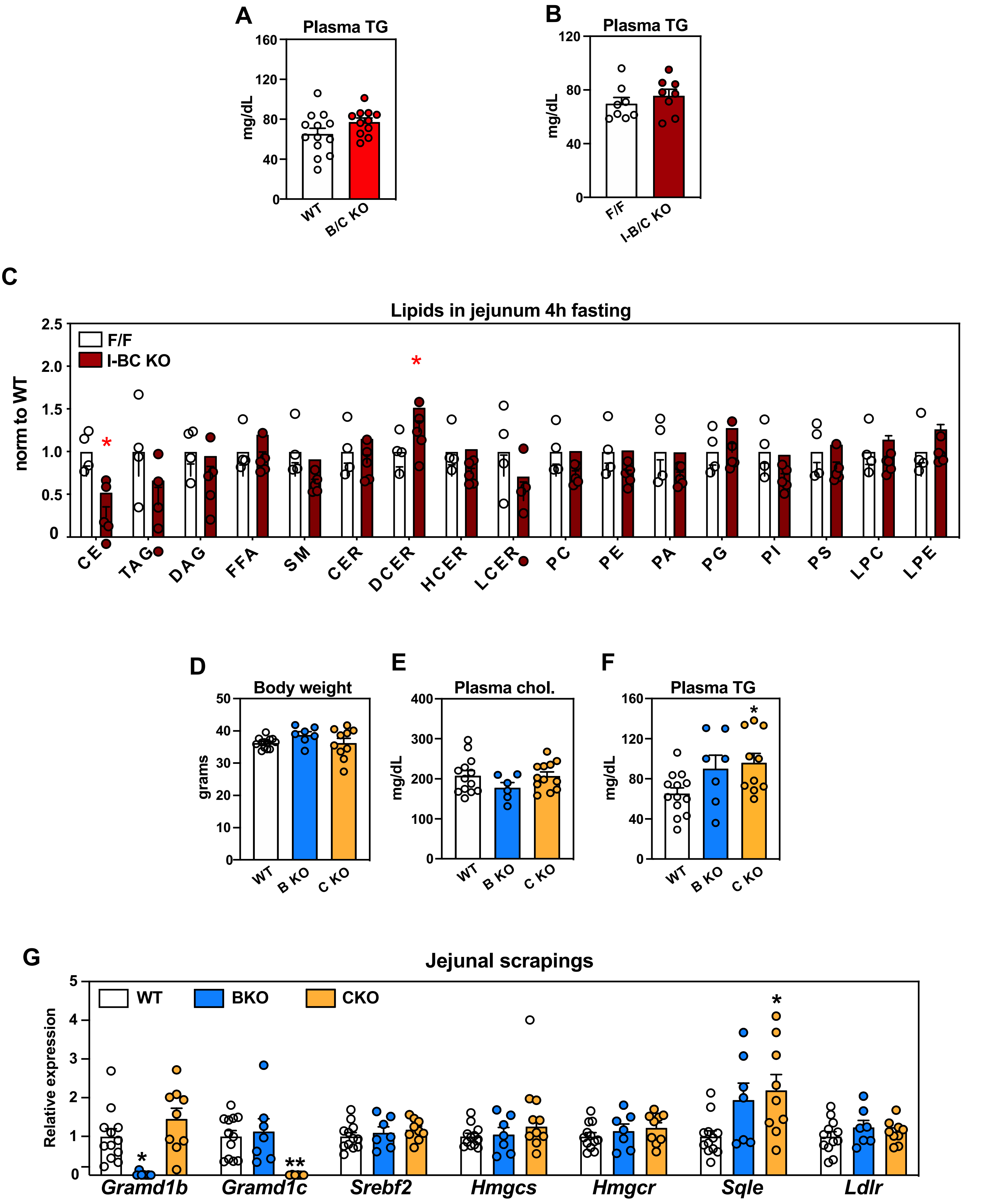


Ferrari et al, Fig. S7

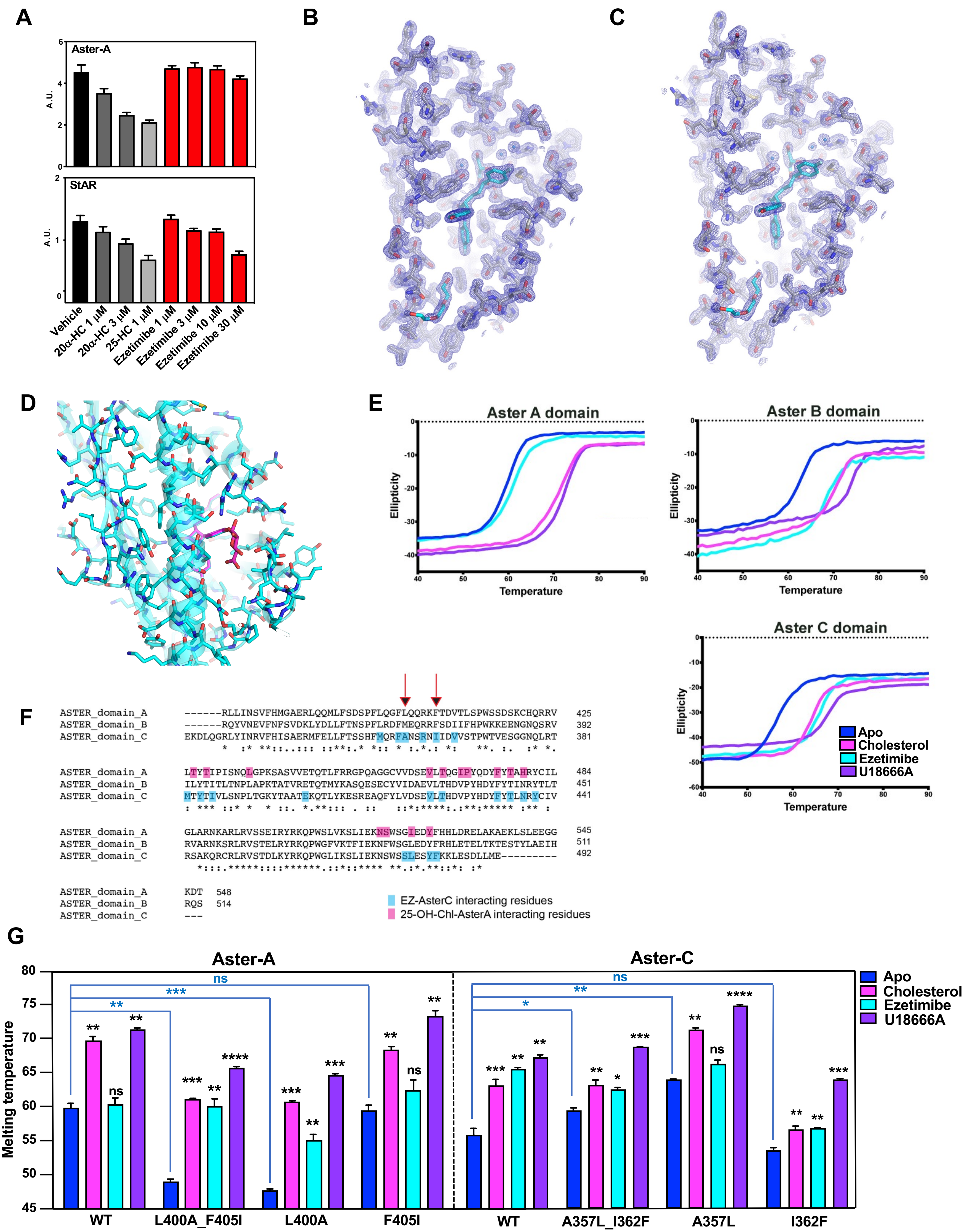


Ferrari et al, Fig. S8

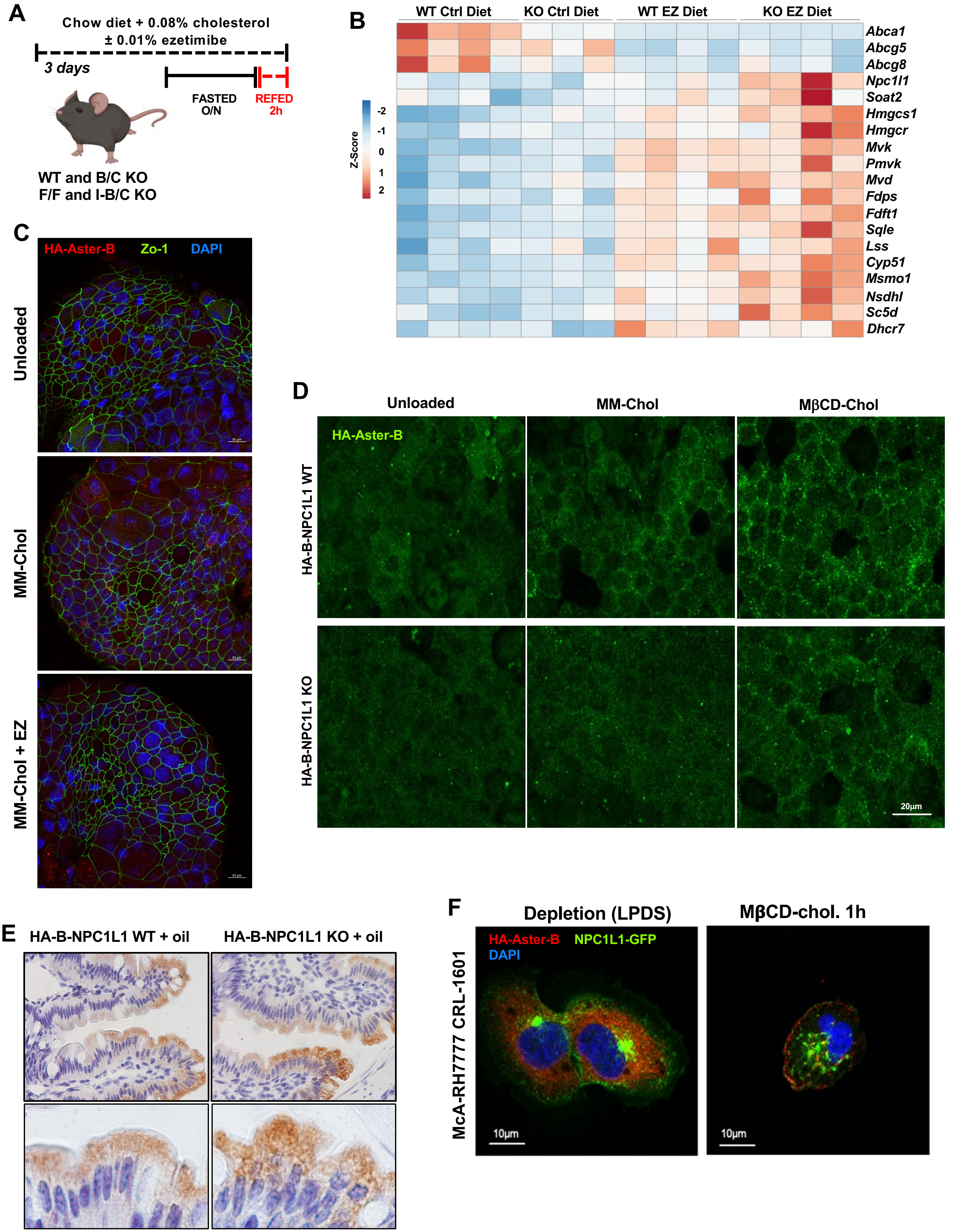


Ferrari et al, Fig. S9

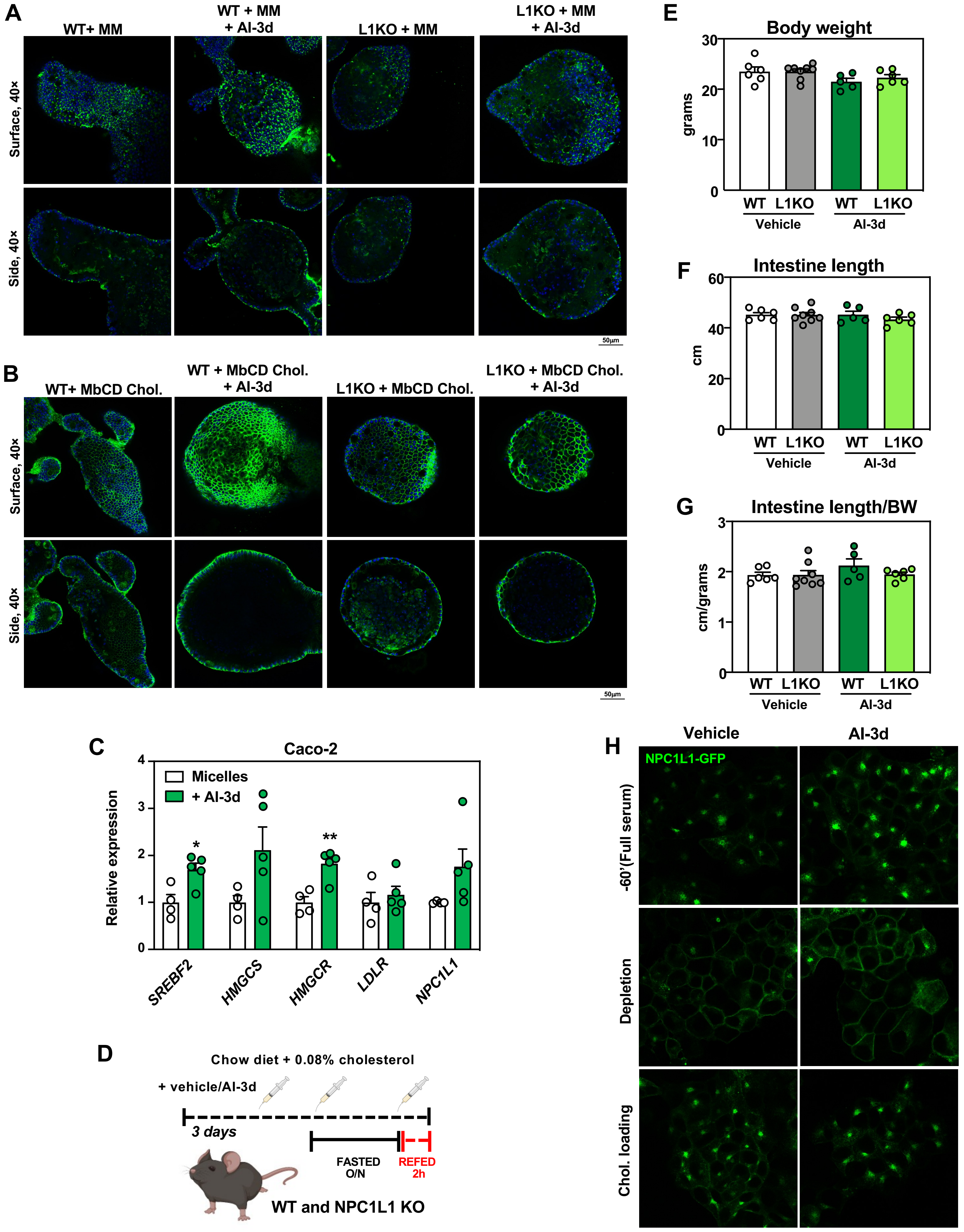


Ferrari et al, Fig. S10
